# The mycobacterial mutasome: composition and recruitment in live cells

**DOI:** 10.1101/2021.11.16.468908

**Authors:** Sophia Gessner, Zela Martin, Michael A. Reiche, Joana A. Santos, Neeraj Dhar, Ryan Dinkele, Timothy De Wet, Atondaho Ramudzuli, Saber Anoosheh, Dirk M. Lang, Jesse Aaron, Teng-Leong Chew, Jennifer Herrmann, Rolf Müller, John D. McKinney, Roger Woodgate, Valerie Mizrahi, Meindert H. Lamers, Digby F. Warner

**Author notes:** These authors contributed equally (Sophia Gessner, Zela Martin, Michael A. Reiche). Vaccine and Infectious Disease Organization (VIDO), University of Saskatchewan, 120 Veterinary Road, Saskatoon, SK, S7N 5E3, Canada. Department of Chemistry and Umeå Centre for Microbial Research, Umeå University, 90187, Umeå, Sweden (Saber Anoosheh).

## Abstract

A DNA damage-inducible mutagenic gene cassette has been implicated in the emergence of drug resistance in *Mycobacterium tuberculosis* during anti-tuberculosis (TB) chemotherapy. However, the molecular composition and operation of the encoded “mycobacterial mutasome” – minimally comprising DnaE2 polymerase and ImuA′ and ImuB accessory proteins – remain elusive. Following exposure of mycobacteria to DNA damaging agents, we observe that DnaE2 and ImuB co-localize with the DNA polymerase III β subunit (β clamp) in distinct intracellular foci. Notably, genetic inactivation of the mutasome in an *imuB*^AAAAGG^ mutant containing a disrupted β clamp-binding motif abolishes ImuB-β clamp focus formation, a phenotype recapitulated pharmacologically by treating bacilli with griselimycin and in biochemical assays in which this β clamp-binding antibiotic collapses pre-formed ImuB-β clamp complexes. These observations establish the essentiality of the ImuB-β clamp interaction for mutagenic DNA repair in mycobacteria, identifying the mutasome as target for adjunctive therapeutics designed to protect anti-TB drugs against emerging resistance.

## INTRODUCTION

*Mycobacterium tuberculosis*, the causative agent of tuberculosis (TB), consistently ranks among the leading infectious killers worldwide (WHO, 2021). The heavy burden imposed by TB on global public health is exacerbated by the emergence and spread of drug-resistant (DR) *M. tuberculosis* strains, with estimates indicating that DR-TB now accounts for approximately one-third of all deaths owing to antimicrobial resistance (Hasan 2018). In the absence of a wholly protective vaccine, a continually replenishing pipeline of novel chemotherapeutics is required (Evans and Mizrahi 2018) which, given the realities of modern antibiotic development (Nielsen 2019), appears unsustainable. Therefore, alternative approaches must be explored including the identification of effective multidrug combinations (Cokol *et al*., 2017), the elucidation of “resistance-proof” compounds (Ling 2015), and the identification of so-called “anti-evolution” drugs that might limit the development of drug resistance (Smith and Romesberg, 2007; Ragheb *et al*., 2019; Merrikh and Kohli, 2020).

Whereas many bacterial pathogens accelerate their evolution by sampling the immediate environment – for example, *via* fratricide, natural competence, or conjugation (von Wintersdorff *et al*., 2016; Veening and Blokesch, 2017) – these mechanisms appear inaccessible to *M. tuberculosis*: the bacillus does not possess plasmids (Gray and Derbyshire, 2018) and there appears to be no role for horizontal gene transfer in the modern evolution of strains of the *M. tuberculosis* complex (Galagan, 2014; Boritsch and Brosch, 2016). Instead, genetic variation in *M. tuberculosis* results exclusively from chromosomal rearrangements and mutations, a feature reflecting its ecological isolation (an obligate pathogen, *M. tuberculosis* has no known host outside humans) and the natural bottlenecks that occur during transmission (Gagneux, 2018). A question which therefore arises is whether a specific molecular mechanism(s) drives *M. tuberculosis* mutagenesis – perhaps under stressful conditions – and, consequently, if the activity thereof might be inhibited pharmacologically.

Multiple studies have investigated mycobacterial DNA replication and repair function in TB infection models (for recent reviews, Singh, 2017; Minias *et al*., 2018; Mittal *et al*., 2020). From these, the C-family DNA polymerase, DnaE2, has emerged as major contributor to mutagenesis under antibiotic treatment (Boshoff *et al*., 2003). A non-essential homolog of *E. coli* DNA Polymerase (Pol) IIIa (Timinskas *et al*., 2014), DnaE2 does not operate alone: the so-called “accessory factors”, *imuA′* and *imuB*, are critical for DnaE2-dependent mutagenesis (Warner *et al*., 2010) (***Figure 1A***). Both proteins are of unknown function, however *imuA′* and *imuB* are upregulated together with *dnaE2* following exposure of mycobacteria to DNA damaging agents including mitomycin C (MMC) (***Figure 1B***). This observation prompted the proposal that these three proteins might represent a “mycobacterial mutasome” – named according to its functional analogy with the *E. coli* DNA Pol V mutasome comprising UmuD′_2_C-RecA-ATP (Jiang *et al*., 2009; Erdem *et al*., 2014).

**Figure 1.**
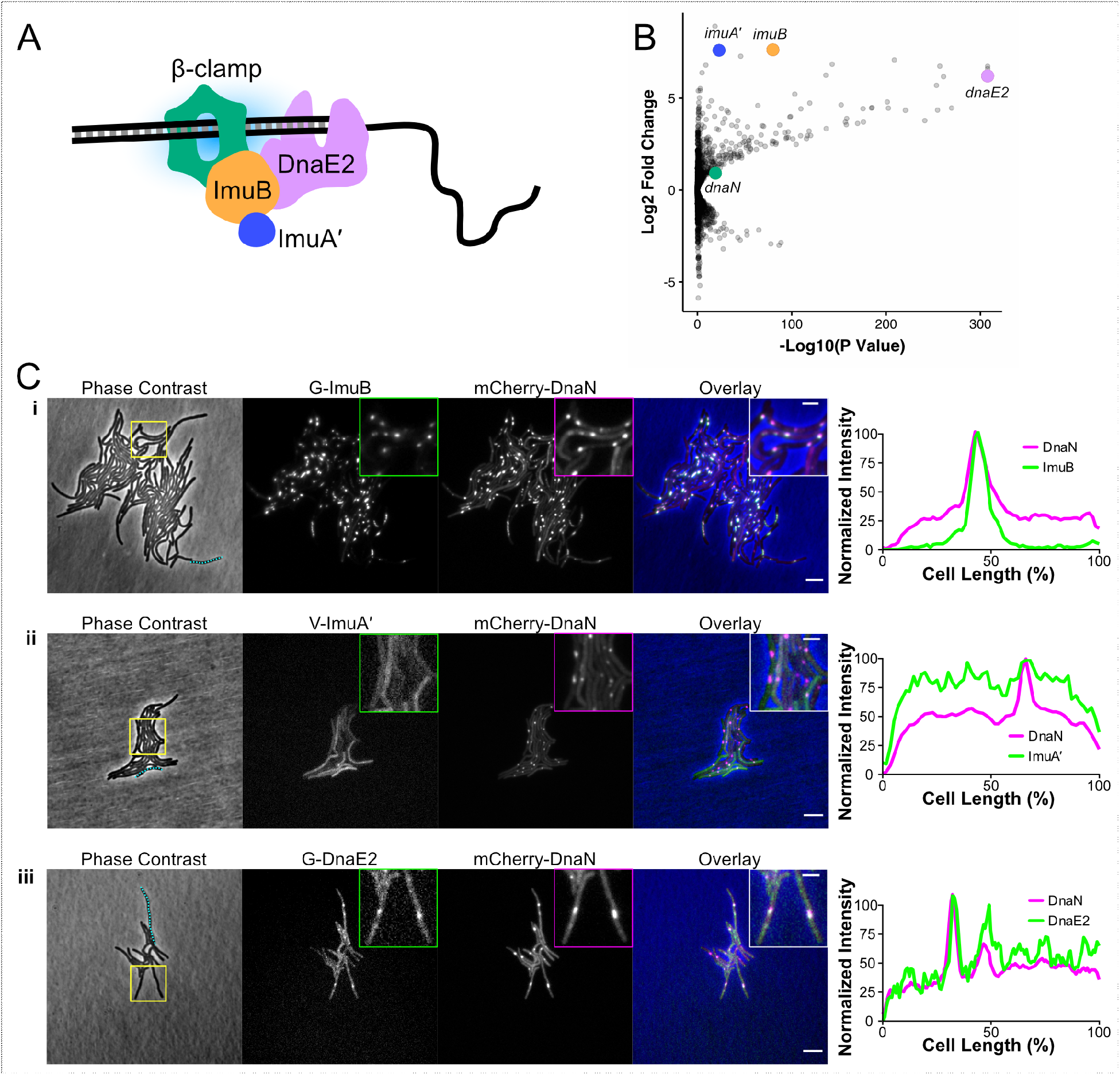
Components of the mycobacterial mutasome. **A.** Cartoon summarizing the available genetic, microbiological and bioinformatic data (Boshoff et al., 2003; Warner et al., 2010; Timinskas and Venclovas, 2019) into a working model of predicted mutasome composition. This model proposes that ImuB (blue) functions as an adapter protein, binding the dnaN-encoded β clamp (orange) to enable access of the error-prone DnaE2 subunit (magenta) to the DNA replication fork. ImuB has also been predicted to bind accessory protein ImuA′ (yellow). **B.** A volcano plot depicting transcriptional data from Müller et al. (2018) which were derived from RNA-seq analyses of wild-type M. smegmatis mc^2^155 exposed to 1×MIC Mitomycin C (MMC). Consistent with corresponding observations in M. tuberculosis (Boshoff et al., 2003; Warner et al., 2010), the mutasome components, imuA′, imuB, and dnaE2 are among the 15 most highly upregulated mycobacterial genes following MMC exposure. **C.** Representative stills from single-cell time-lapse fluorescence microscopy experiments of M. smegmatis expressing translational reporters of the different mutasome components. Phase contrast and fluorescence images of M. smegmatis expressing G-ImuB and mCherry-DnaN, V-ImuA′and mCherry-DnaN, and G-DnaE2 and mCherry-DnaN are represented following 4 h exposure to ½×MIC MMC. Scale bars in the overlay image represent 5 μm. Inset images at the top right corner of the fluorescence and overlay images show a zoomed-in region corresponding to the yellow box in each phase contrast image (inset scale bar represents 2 μm). The right-hand panel illustrates normalised fluorescence intensity along the longitudinal axis (as percentage [%] of the total cell length) of a representative cell from each strain; the cell analysed is indicated by the blue dotted lines in the corresponding phase contrast image. Single-cell time-lapse fluorescence microscopy experiments were repeated 2-4 times; a typical experiment collected images from approximately 80 XY points.

Here, we apply live-cell fluorescence and time-lapse microscopy in characterizing a panel of mycobacterial reporter strains expressing fluorescent translational fusions of each of the known mutasome components. The results of these analyses, together with complementary *in vitro* biochemical assays utilizing purified mycobacterial proteins, support the inference that ImuB serves as a hub protein, interacting with the *dnaN*-encoded β clamp and ImuA′. They also reinforce the essentiality of the ImuB-β clamp protein-protein interaction for mutasome function. Notably, while a strong ImuA′-ImuB interaction is detected *in vitro*, we report live-cell data which indicate the dispensability of either ImuA′ or DnaE2 for ImuB localization – but not mutasome function – in bacilli exposed to genotoxic stress. Finally, using the β clamp-binding antibiotic, griselimycin (GRS) (Kling *et al*., 2015), we demonstrate in biochemical assays and in live mycobacteria the capacity to inhibit mutasome function through the pharmacological disruption of ImuB-β focus formation. These observations suggest that, through its inhibition of β clamp binding, GRS naturally limits the capacity for induced mutagenesis. Therefore, as well as revealing a built-in mechanism protecting against auto-induced mutations to GRS resistance, our results support the potential utility of “anti-evolution” antibiotics for TB.

## RESULTS

### ImuB forms distinct sub-cellular foci under DNA damaging conditions

Our previous genetic evidence (Warner *et al*., 2010) informed a tentative model in which the catalytically inactive Y family Pol homolog, ImuB, functioned as an adapter protein. According to the model, DnaE2 gains access to the repair site by interacting with ImuB, which similarly interacts with ImuA′ and the dnaN-encoded β clamp subunit (***Figure 1A***). To investigate the subcellular localizations of each of the mutasome proteins in live bacilli, we constructed reporter alleles in which the *M. smegmatis* mutasome proteins were labelled by N-terminal translational attachment of either Enhanced Green (EGFP) or Venus Fluorescent Protein (VFP) tags (***Figure 1 – figure supplement 1A***). The reporter alleles were introduced into each of three individual *M. smegmatis* mutasome gene deletion mutants – Δ*dnaE2*, Δ*imuA′*, and Δ*imuB* (Warner *et al*., 2010) – to yield the complemented strains, Δ*dnaE2 attB*::*egfp-dnaE2* (designated G-*dnaE2*), Δ*imuB attB*::*egfp-imuB* (G-*imuB*), and Δ*imuA′attB*::*vfp-imuA′* (V-*imuA′*).

The mycobacterial DNA damage response was induced by exposing the strains to the natural product antibiotic, mitomycin C (MMC), an alkylating agent that causes monofunctional DNA adducts and inter-strand and intra-strand cross-links (Bargonetti *et al*., 2010). Following exposure of V-*imuA′* to MMC for 4 hours, a yellow fluorescence signal was observable throughout the cells (***Figure 1C, Figure 1 – figure supplement 1B***), suggesting diffuse distribution of the VFP-ImuA′ protein in the mycobacterial cytoplasm. In contrast, distinct EGFP-ImuB foci were observed in G-*imuB* bacilli following MMC treatment (***Figure 1C, Figure 1 – figure supplement 1B***). Although less distinct, EGFP-DnaE2 produced similar evidence of focus formation in G-*dnaE2* cells (***Figure 1C, Figure 1 – figure supplement 1B***). Notably, the significant increase in signal detectable in V-*imuA′*, G-*imuB*, and G-*dnaE2* for MMC-exposed *versus* unexposed cells (***Figure 1 – figure supplement 1B***) confirmed that expression of the respective fluorescence reporter alleles was DNA damage-dependent in all three reporter mutants.

To determine whether these observations were also true for other types of DNA damage, the three reporter mutants were subjected to ultra-violet (UV) light exposure. Equivalent fluorescence phenotypes were observed for each of the three reporter alleles under both DNA damaging treatments (***Figure 1 – figure supplement 1B)***. As UV exposure causes cyclobutane pyrimidine dimers or pyrimidine-pyrimidone (6-4) photoproducts (Boshoff *et al*., 2003), while MMC generates inter-strand DNA cross-links at CpG sites (Tomasz, 1995), these results indicated that expression and localization (recruitment) of the mutasome components might be independent of the nature of the genotoxic stress applied.

### N-terminal fluorescent reporters retain wild-type mutagenic function but are deficient in DNA damage tolerance

The addition of bulky fluorescent tags can disrupt the function of DNA replication and repair proteins (Renzette *et al*., 2005). To determine whether any of the tagged mutasome proteins were affected, the functionalities of the *egfp-imuB, vfp-imuA′* and *egfp-dnaE2* alleles were assessed in two standard assays (Boshoff *et al*., 2003; Warner *et al*., 2010): the first investigated DNA damage-induced mutagenesis following UV irradiation, and the second tested DNA damage tolerance following treatment with MMC. As observed previously (Boshoff *et al*., 2003; Warner *et al*., 2010), exposure of the wild-type parental *M. smegmatis* mc^2^155 to a sub-lethal dose of UV irradiation increased the frequency of rifampicin (RIF) resistance 50-to 100-fold, as determined from enumeration of colony forming units (CFU) on RIF-containing solid growth medium. In contrast, induced mutagenesis was greatly reduced in the Δ*imuA′, DimuB*, and *DdnaE2* deletion mutants: mutation frequencies for these “mutasome-deficient” strains were approximately 20-fold lower than wild-type (***Figure 1 – figure supplement 2A***). Complementation with the cognate fluorescent reporter allele in V-*imuA′*, G-*imuB* and G-*dnaE2* restored the UV-induced mutation frequency of each of the three knockout mutants to near wild-type levels, establishing that each of the fluorescence reporter alleles retained function in UV-induced mutagenesis assays.

Surprisingly, the DNA damage tolerance assay – in which CFU forming ability is tested during continuous exposure to MMC in solid medium – produced contrasting results (***Figure 1 – figure supplement 2B***): unlike in the mutagenesis assay, complementation of either *imuA’* or *imuB* gene deletion mutants with their corresponding fluorescent reporter alleles failed to restore the wild-type phenotype, whereas *dnaE2* hypersusceptibility was reversed. The reason for these discrepant observations – restoration of UV-induced mutagenesis but not MMC-induced DNA damage tolerance – in the *imuA* and *imuB* strains is not clear. It seems likely that the different types of DNA damage induced by the two separate treatments might require distinct repair pathways and, potentially, discrete protein interactions which were differentially disrupted by the presence of a fluorescent tag on either mutasome component. Another possibility is that this phenotype was caused by the persistent/recurring damage sustained by the bacilli throughout the 4-day incubation on MMC-containing medium – dissimilar to the comparatively short duration of UV exposure. However, these explanations are speculative and require further investigation.

### ImuB localizes with the *dnaN*-encoded β clamp following DNA damage

We previously inferred that a putative interaction between ImuB and the *dnaN*-encoded β clamp was essential for mutasome function (Warner *et al*., 2010). To investigate the predicted interaction of ImuB and the β clamp in live bacilli, each of the three mutasome reporter alleles was introduced separately into an *M. smegmatis* mutant encoding an mCherry-tagged β clamp, mCherry-DnaN (Santi *et al*., 2013). The mCherry-DnaN reporter was chosen as background strain owing to its previous validation in single-cell time-lapse fluorescence microscopy analyses of *M. smegmatis* replisome location (Santi *et al*., 2013; Santi and McKinney, 2015). For the time-lapse experiments, the *M. smegmatis* dual reporter strains were grown in standard 7H9/OADC medium for 12 hours, following which the cells were exposed to MMC for 4.5 hours before switching back to 7H9/OADC for post-treatment recovery (***Figure 1C**; **Figure 2**; **Videos 1-3***). At 4 hours post MMC treatment, distinct EGFP-ImuB foci were observed (***Figure 1C panel i***) which, when overlaid with the mCherry-DnaN fluorescence signal, showed considerable overlap, suggesting association of the β clamp with ImuB. Notably, the EGFP-ImuB signal was almost exclusively detected in very close proximity to mCherry-DnaN foci, with very rare instances of EGFP signal detectable in regions where fluorescence was absent. This association was independent of the duration of MMC exposure, occurring at all time points tested (***Figure 2A; Video 1***). In combination, these results support the direct physical interaction of ImuB and the β clamp, as suggested previously by yeast two-hybrid and site-directed mutagenesis studies (Warner *et al*., 2010).

**Figure 2.**
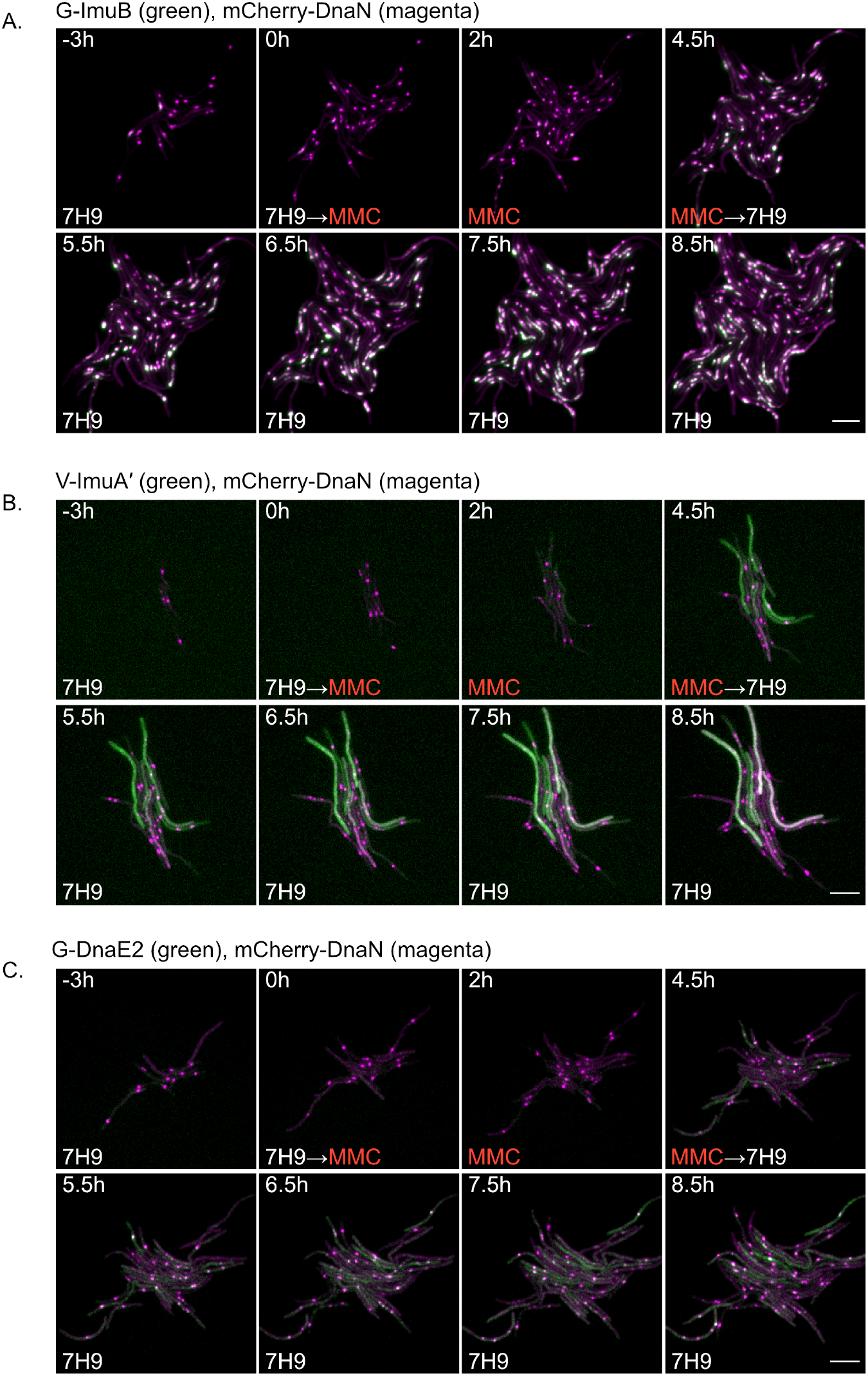
Representative time-lapse series of single-cells of M. smegmatis expressing the generated translational reporters in combination with mCherry-DnaN. **A.** G-ImuB (green) and mCherry-DnaN (magenta), **B.** V-ImuA′ (green) and mCherry-DnaN (magenta), and **C.** G-DnaE2 (green) and mCherry-DnaN (magenta). Overlapping signals are viewed as white. The cells were exposed to MMC (½×MIC), from time 0 h until 4.5 h, after which the medium was switched back to standard 7H9 OADC medium. Up to 80 XY points were imaged at 10-minute intervals on fluorescence and phase channels for up to 36 h. The experiments were repeated 2–4 times. Numbers indicate hours elapsed. Scale bar represents 5μm. 7H9: Middlebrook 7H9 medium. MMC: Mitomycin C.

For V-ImuA′, a diffuse fluorescence signal was detected throughout the cells (***Figure 1C panel ii; Figure 2B; Video 2***), rendering impossible any conclusion about the potential recruitment of ImuA′ to β clamp (mCherry-DnaN) foci. In contrast, the results for DnaE2 were more nuanced: overlap of peak fluorescence signals from EGFP-DnaE2 and mCherry-DnaN proteins was detected (***Figure 1C panel iii; Figure 2C***) and was most evident within 1 hour post removal of MMC from the microfluidic chamber (***Video 3***). Although not as consistent as the ImuB-β clamp phenotype, the co-localization was reproducibly observed in multiple cells and across different experiments.

### ImuA′ and DnaE2 are not required for ImuB focus formation

We showed previously that deletion of imuA′ phenocopied abrogation of either imuB or dnaE2 in the MMC sensitivity assay (Warner, *et al*., 2010) and, consistent with the interpretation that all three components are individually essential for mutasome activity, this phenotype was not exacerbated in a triple knockout mutant (Δ*imuA′-imuB* Δ*dnaE2*) lacking all three mutasome components. Moreover, yeast two-hybrid results predicted a direct interaction between ImuB and ImuA′ (Warner *et al*., 2010). Together, these observations suggested that a deficiency in ImuA′ might impair ImuB protein localization. To test this prediction, the *egfp-imuB* allele was introduced into the ΔimuA′ deletion mutant, generating a Δ*imuA′ attB*::*egfp-imuB* reporter strain. Despite the loss of ImuA′ in this mutant, EGFP-ImuB foci were observed following MMC treatment (***Figure 2 – figure supplement 1i***), mimicking the wild-type phenotype. The absence of functional DnaE2 similarly had no discernible impact on ImuB focus formation in the corresponding catalytically dead *dnaE2*^AIA^ *attB*::*egfp-imuB* or DnaE2-deleted Δ*dnaE2 attB*::*egfp-imuB*, mutants (***Figure 2 – figure supplement 1ii & iii***). In combination, these results appear to eliminate a role for either ImuA′ or DnaE2 in ImuB localization, instead implying the critical importance of the ImuB-β clamp interaction for mutasome function.

### Purified mutasome proteins interact in biochemical assays *in vitro*

All inference from this and previous work about the composition of the mycobacterial mutasome has been derived from microbiological assays. To address this limitation, we expressed and purified recombinant *M. smegmatis* mutasome proteins for biochemical analysis. Expression in *E. coli* of ImuB alone yielded low quantities of soluble protein that was prone to degradation, while attempts to express ImuA′alone failed to generate soluble protein. In contrast, co-expression of ImuB with ImuA′yielded both proteins in a soluble form (***Figure 3***). Subsequently, the ImuA′B complex could be captured *via* a histidine (His) affinity tag in ImuB. This confirmed that ImuA′and ImuB interact *in vitro*, forming a stable complex even at protein concentrations as low as 400 nM (***Figure 3 – figure supplement 1***), corroborating previous yeast two-hybrid results (Warner *et al*., 2010). In *E. coli*, overexpression of DnaE2 resulted in insoluble protein, while DnaE2 overexpression in *M. smegmatis* appeared to be incompatible with cell viability: following transformation with the expression construct, very few colonies were obtained and could not be expanded in liquid culture (not shown).

**Figure 3.**
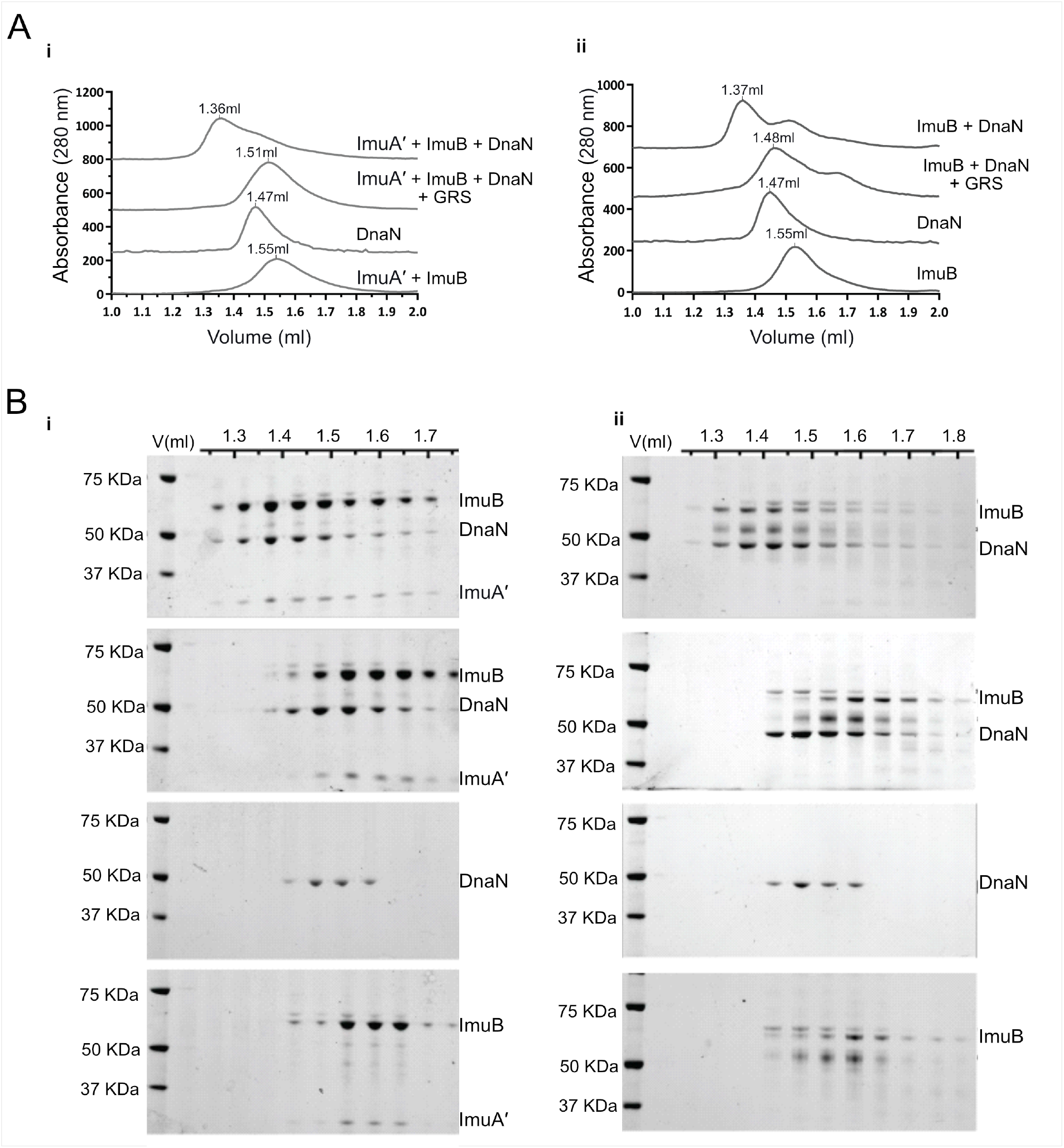
ImuB and ImuA′-ImuB interact with the DnaN and these interactions are disrupted by GRS. **A.** Gel filtration profiles of M. smegmatis (i) ImuA′-ImuB-DnaN and (ii) ImuB-DnaN complexes in the absence or presence of 15 μM GRS. For these experiments, 5 μM DnaN was added to 10 μM of (i) ImuA′-ImuB or (ii) ImuB. The gel filtration profiles of the individual proteins (ImuB and DnaN) or complex (ImuA′-ImuB) are shown for comparative purposes, and all curves were scaled for clarity. **B.** SDS–PAGE analysis of sequential fractions of the gel filtration runs. Gels are sorted in the same order as the corresponding gel filtration profiles shown in A.

Next, we analysed the interaction of the *dnaN*-encoded β clamp with ImuB or the ImuA′B complex (***Figure 3***). Samples of the *M. smegmatis* β clamp with ImuA′B (***Figure 3A panel i***) or ImuB (***Figure 3A panel ii***), were injected onto an analytical size-exclusion chromatography column and collected fractions were analysed by SDS-PAGE. Alone, the β clamp and ImuB/ImuA′B eluted at 1.47 and 1.55 ml, respectively. Incubation of the β clamp with either ImuB or ImuA′B caused a shift in the retention volume to 1.36 ml, indicative of complex formation. This was confirmed by SDS-PAGE analysis, which indicated co-elution of the β clamp with ImuB and ImuA′B.

### GFP-ImuB and VFP-ImuA′ form a stable complex

Our microbiological assays had unexpectedly revealed discrepant complementation phenotypes for UV exposure (***Figure 1 – figure supplement 2A***) versus MMC treatment (***Figure 1 – figure supplement 2B***), raising the possibility that the fluorescent tags in the bioreporter mutants might disrupt a protein-protein interaction(s) essential for DNA damage tolerance. We therefore investigated the capacity of the fluorescently labelled EGFP-ImuB and VFP-ImuA′ proteins to form a stable complex. To this end, His-EGFP-ImuB was co-expressed with Strep-VFP-ImuA′ in *E. coli* and the complex analysed in three consecutive chromatography steps (***Figure 3 – figure supplement 1B***). First, the cell lysate was loaded onto a HisTrap column to capture the VFP-ImuA′:EGFP-ImuB complex via the His-tag present in EGFP-ImuB. Next, the elution fractions containing the complex were loaded on a StrepTrap column to capture the complex via the strep-tag on VFP-ImuA′. Finally, the VFP-ImuA′:EGFP-ImuB complex was injected onto a size-exclusion column.

During all purification steps, EGFP-ImuB with VFP-ImuA′ were co-eluted as a complex, as indicated by SDS-PAGE analysis and fluorescent detection of GFP-ImuB and VFP-ImuA′ in the same elution fractions. In combination, these observations suggest that the fluorescent tags did not disrupt ImuA′-ImuB complex formation – a result which additionally implies that the absence in live cells of a clear ImuA′ (co-)localization phenotype was not a function of the tags.

### Inhibition of ImuB-β clamp binding eliminates focus formation

Previous work established that the β clamp-binding domain of ImuB was essential for mutasome function: mutant strains carrying either a *imuB*^Δ_C168_^ allele – which lacks the 168 amino acids in the ImuB C-terminal region – or a *imuB*^AAAAGG^ allele – in which the wild-type β clamp-binding motif, ^352^QLPLWG^357^, is substituted with the non-functional ^352^AAAAGG^357^ peptide sequence – phenocopied full *imuB* deletion (Warner *et al*., 2010). Therefore, to test the prediction that the SOS-regulated recruitment of EGFP-ImuB and mCherry-β clamp into discernible foci was dependent on the ImuB-β protein-protein interaction, we introduced an *egfp-imuB*^AAAAGG^ allele into the Δ*imuB* mutant. In contrast to the wild-type reporter (G-*imuB*), the β clamp-binding motif mutant (G-*imuB*^AAAAGG^) exhibited no EGFP foci in any cell imaged following exposure to MMC (***Figure 4A panel ii***). Instead, the fluorescence was detectable throughout the cell as a diffuse signal (***Figure 4B***). This result supports the inferred essentiality of the physical interaction between ImuB and β for ImuB localization and, moreover, establishes that detection of ImuB-β foci provides a reliable visual proxy for functional mutasome formation.

**Figure 4.**
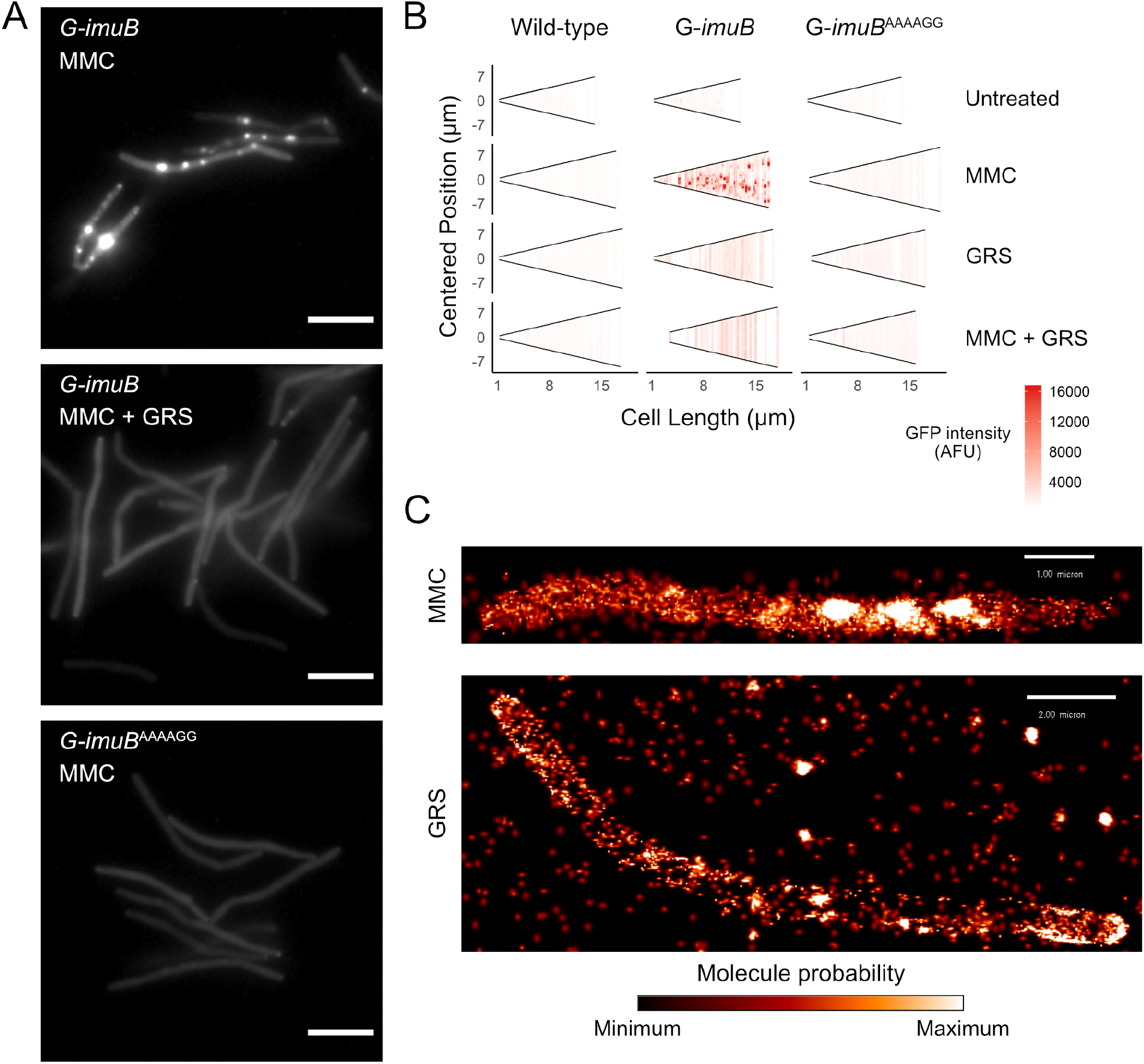
Disrupting the DnaN-ImuB interaction. **A.** Representative images of G-imuB exposed to 2× MIC MMC for 4 hr (top panel) or 2× MIC MMC and GRS for 4 hr (centre panel) and G-imuB^AAAAGG^ mutant exposed to 2× MIC MMC for 4 hr (bottom panel). Scale bars represent 5 μm. **B.** Cells aligned by mid-cell position, arranged according to cell length and coloured (white to red) according to fluorescence intensity showing the presence of G-ImuB foci following MMC treatment and the lack thereof after GRS treatment. G-imuB^AAAAGG^ shows no foci, similar to the G-imuB strain following GRS exposure. **C.** Super-resolution imaging confirms disruption of ImuB-β clamp foci in mycobacteria treated with GRS. Representative iPALM localizations of PC-ImuB bacilli exposed to (top) 5× MIC MMC or (bottom) 5× MIC GRS. Subdiffraction-limited super-resolution localization of PC-ImuB is observed as highly dense localizations of signal following exposure to MMC; in contrast, GRS prevents the formation of high-density localizations. Scale bars are 1 μm and 2 μm in the top and bottom micrographs, respectively; molecule probability represents the fluorescence localization probability from minimum (black) to maximum (white) likelihood.

### Griselimycin blocks ImuB-β clamp binding, preventing focus formation in *M. smegmatis*

Griselimycin (GRS) is a natural product antibiotic that binds the mycobacterial β clamp with high affinity, preventing DNA replication by blocking the essential interaction with the PolIIIa subunit, DnaE1 (Kling *et al*., 2015). Notably, the location of GRS binding overlaps with the region predicted to interact with other β clamp-binding proteins, including ImuB (Bunting *et al*., 2003; Burnouf *et al*.,2004; Kling *et al*., 2015). Therefore, we hypothesized that GRS might disrupt the ImuB-β interaction. Indeed, addition of GRS disrupted the *in vitro* interaction between the β clamp and pre-formed ImuA′B complex (***Figure 3A panel i***) as well as between the β clamp and ImuB (***Figure 3A panel ii***), as indicated by a gel filtration profile that is a superposition of the absorbance traces of the sample individual components (β clamp and ImuB or β clamp and ImuA′B). This was confirmed by SDS-PAGE analysis (***Figure 3B***). To confirm that the disrupting effect of GRS on the complex was the result of the GRS-β clamp binding (Kling *et al*., 2015), we measured the melting curves of the β clamp in the presence and absence of GRS (***Figure 3 – figure supplement 1C***). Incubation with GRS led to a 3 °C increase in the protein melting temperature, consistent with GRS binding to β. In contrast, no effect of GRS was observed on the melting temperatures of ImuA′B.

Next, we examined whether these *in vitro* observations were recapitulated in live mycobacterial cells. To this end, the G-*imuB* strain was exposed to MMC (***Figure 4A panel i***) or MMC plus GRS (***Figure 4A panel iii***). Notably, the addition of GRS in combination with MMC abrogated G-*imuB* focus formation, phenocopying the diffuse fluorescence distribution observed following exposure of the β clamp-binding deficient GFP-ImuB^AAAAGG^ mutant to MMC ***Figure 4A panel ii***). Population analyses of these data confirmed that GRS blocked ImuB focus formation in MMC-exposed cells (***Figure 4B***), suggesting the potential for chemical disruption of mutasome function.

### Super-resolution microscopy confirms GRS elimination of ImuB-β foci in *M. smegmatis*

Finally, to examine the inferred disruption of ImuB focus formation by single-molecule microscopy, we generated an additional reporter mutant in which ImuB was N-terminally labeled with the photoconvertible, fixation-resistant mEos4a fluorophore (Paez Segala *et al*., 2015). The resulting PC-*imuB* strain was imaged in 3D by iPALM (Shtengel *et al*., 2009) following exposure to MMC or GRS (***Figure 4C***). Consistent with the epifluorescence data, bacilli exposed to MMC were characterized by a region of high-density PC-ImuB localizations (***Figure 4C top panel***). In contrast, exposure to GRS – which on its own has been shown to induce the *M. smegmatis* SOS response (Kling *et al*., 2015) – did not elicit high molecule densities: instead, a low-density of molecules was detected throughout the interior of the cell, reinforcing the inferred absence of ImuB recruitment in GRS-exposed cells (***Figure 4C bottom panel***).

## DISCUSSION

In *E. coli*, the DNA damage-induced SOS response triggers overexpression of *umuC, umuD* and *recA* (Maslowska *et al*., 2019). UmuC is an error prone Y-family DNA polymerase that requires the binding of UmuD′_2_ and RecA to reach full activity and this “mutasome”, collectively referred to as PolV, has been implicated in DNA damage tolerance and induced mutagenesis (Goodman *et al*., 2016). At the onset of the work reported here, genetic evidence from diverse bacteria lacking PolV supported the co-dependent operation of ImuA, ImuB, and DnaE2 in the LexA-regulated SOS response, suggesting these proteins might function in an analogous manner (McHenry, 2011; Ippoliti *et al*., 2012). In mycobacteria, in which they have also been implicated in DNA damage tolerance and induced mutagenesis (Boshoff and Mizrahi, 2000; Warner *et al*., 2010), the non-homologous ImuA′ replaces ImuA, nevertheless the inferred universal model for mutasome function in bacteria lacking an *E. coli* PolV homolog was the same (Timinskas and Venclovas, 2019): the catalytically inactive Y family polymerase, ImuB, functions as hub protein, interacting physically with the β clamp via a defined β clamp-binding motif and with DnaE2 and ImuA′ (or ImuA) via unknown mechanisms which might include the disordered ImuB C-terminal region or sub-regions thereof, including the RecA-NT motif (Timinskas and Venclovas, 2019). However, the absence of any direct biochemical and/or structural evidence to support the proposed protein interactions meant this assumption was speculative. Moreover, whereas *E. coli* PolV is known to be subject to multiple forms of regulation – including temporal (Robinson *et al*., 2015), spatial (Robinson *et al*., 2015), internal (Erdem *et al*., 2014) and conformational (Jiang *et al*., 2009; Gruber *et al*., 2015; Jaszczur *et al*., 2019) – the expression dynamics and sub-cellular localizations of the mycobacterial mutasome proteins were mostly unknown.

By fluorescently tagging the known mutasome proteins, we have observed in real-time the consistent formation of co-localizing ImuB-β clamp foci in mycobacterial cell populations exposed to genotoxic stress. Although less pronounced than ImuB, we also detected the frequent, reproducible co-localization of DnaE2 with the β clamp under the same conditions. Notably, recruitment of ImuB into foci occurred in mutants lacking either *dnaE2* or *imuA′* but was prevented where the ImuB β clamp-binding motif was mutated – apparently identifying the primacy of the ImuB-β clamp interaction in mutasome organization. In contrast, the function(s) and sub-cellular dynamics of ImuA′ remain enigmatic: VFP-ImuA′ consistently produced diffuse fluorescence in DNA-damaged bacilli, precluding any definitive insights into its potential association with ImuB (or DnaE2) *in vivo*. We did, however, observe co-elution of ImuA′-ImuB and ImuA′-ImuB-β clamp complexes *in vitro*, results which provided important biochemical confirmation of the inferred interaction of ImuA′ and ImuB predicted previously (Warner *et al*., 2010). While difficult to reconcile with these data, the absence of a clear co-localization signal in live cells might indicate the transient association of ImuA′ with its mutasome partners, or possibly a modification analogous to the proteolytic cleavage of UmuD to UmuD′ in the *E. coli* SOS response (Goodman *et al*., 2016). Future work will require single-molecule tracking of ImuA′ to resolve this possibility.

The original identification of the *imuA*-*imuB*-*dnaE2* cassette noted its close association with LexA across diverse bacteria; that is, genomes containing the cassette invariably encoded a LexA homolog, too (Erill *et al*., 2006). Recent work in mycobacteria has, however, added unexpected nuance to that regulatory framework, namely that the split *imuA′-imuB/dnaE2* cassette is subject to transcriptional control by both the “classic” LexA/RecA-regulated SOS response and the PafBC-mediated DNA damage response. The authors also report that, while the two regulatory mechanisms are partially redundant for genotoxic stresses including UV and MMC exposure, fluoroquinolones appear to be specific inducers of PafBC only (Adefisayo *et al*., 2021). In addition to suggesting that chromosomal mutagenesis is co-dependent on PafBC and SOS, these observations are important in identifying an apparent “fail-safe” mechanism in mycobacteria in which the mutasome components are induced irrespective of DNA damage type – again reinforcing the centrality of these proteins in damage tolerance and, by implication, adaptive mutagenesis.

We previously observed that the *imuB*^AAAAGG^ β clamp-binding mutation eliminated UV-induced mutagenesis and MMC damage tolerance in *M. smegmatis* (Warner *et al*., 2010), phenocopying deletion of any of the three mutasome components (*imuA′*, *imuB*, *dnaE2*) alone or in combination. Given the abrogation of ImuB focus formation, it is reasonable to infer a direct link between ImuB-β clamp focus formation and mutasome function. In turn, this suggests that blockade of ImuB focus formation might offer a tractable read-out for a screen designed to identify mutasome inhibitors – a possibility reinforced by the observed co-elution in biochemical assays of β with ImuB and, separately, of the β clamp with pre-formed ImuA′-ImuB complexes. In this context, it was notable in the current work that GRS disrupted the ImuB-β clamp interaction *in vitro* and prevented ImuB focus formation in mycobacteria treated simultaneously with MMC and GRS.

The discrepant complementation phenotypes observed for V-*imuA′* and G-*imuB* in the DNA damage tolerance (MMC treatment) *versus* induced mutagenesis (UV exposure) assays suggests that addition of the bulky fluorophore might have prevented full function of these mutasome proteins. Whereas UV irradiation predominantly generates cyclobutane dimers and pyrimidine-pyrimidone (6–4) photoproducts (Franklin *et al*., 1985), MMC induces different DNA lesions, including inter- and intra-strand crosslinks. These are likely to require multiple repair proteins and, potentially, the interaction of mutasome components with additional proteins – which might be prevented by the bulky fluorescent tags. The DnaE2-EGFP fusion proved the exception; in this context, it might be instructive to consider recent evidence implicating DnaE2 in gap filling following nucleotide excision repair in non-replicating *Caulobacter crescentus* cells (Joseph *et al*., 2021). These observations suggest the importance of identifying other potential interacting partners of mycobacterial DnaE2 (and the other mutasome components), work which is currently underway in our laboratory.

The potential for inhibitors of DNA replication to accelerate the development of genetic resistance through the induction of mutagenic repair pathways (Cirz *et al*., 2005; Barrett *et al*., 2019; Revitt-Mills and Robinson, 2020) is a valid and commonly cited concern that might partially explain the relative under-exploration of DNA metabolism as source of new antibacterial drug targets (Reiche *et al*., 2017; van Eijk *et al*., 2017). Our results suggest that GRS could offer an interesting exception: that is, in binding the β clamp at the site of interaction with the DnaE1 replicative DNA polymerase (Kling *et al*., 2015) as well as other DNA metabolizing proteins, including ImuB, GRS appears to possess an intrinsic protective mechanism against induced mutagenesis – blocking both ImuB-dependent mutasome recruitment to stalled replisomes and post-repair fixation of mutations by the replicative polymerase, DnaE1. This “resistance-proofing” capacity, which is supported by the reported restriction of GRS resistance to low-frequency, high-fitness cost amplifications of the *dnaN* genomic region with very few to no “off-target” SNPs, might also contribute to the observed bactericidal effect of GRS against mycobacteria (Kling *et al*.,2015). In addition, it reinforces the β clamp as vulnerable target for new TB drug development (Bosch *et al*., 2021). In this context, it is worth noting that inhibition of DnaE1 replicative polymerase function might represent a general solution to the problem of drug-induced (auto)mutagenesis by preventing fixation of repair/tolerance-generated mutations; in support of this inference, another natural product, nargenicin, which inhibits *Mtb* DnaE1 via a DNA-dependent mechanism, fails to yield spontaneous resistance mutations *in vitro* (Chengalroyen *et al*., 2021). Therefore, while the essentiality of DNA replication proteins (including DnaN, DnaE1) for mycobacterial viability poses a challenge to the design of assays of “anti-evolution” compounds targeting these proteins, GRS (and nargenicin) appear to provide compelling evidence that precise inhibition of specific DNA replicative and repair functions might ameliorate the perceived risks in targeting this area of mycobacterial metabolism.

As an obligate human pathogen, persistence of *M. tuberculosis* within its host depends on the ability to drive successive cycles of infection, disease – in some cases latency followed by reactivation disease – and transmission (Lin and Flynn, 2018). This process is necessarily vulnerable to multiple potential evolutionary culs-de-sac which might arise in consequence of the elimination of the bacillus by the host (clearance) or the demise of the organism within the infected individual (controlled subclinical infection, or host death). Modern *M. tuberculosis* strains therefore represent the genotypes that have successfully adapted to human colonization (Gagneux, 2018), evolving with their obligate host through changes in lifestyle and nutritional habits (with their associated implications for non-communicable diseases such as diabetes), the near-universal administration of the BCG vaccination, the emergence of the HIV co-pandemic, and the widespread use of frontline combination chemotherapy (Warner *et al*., 2015). While the emergence and propagation of drug-resistant isolates characterized by a variety of polymorphisms at multiple genomic loci (Warner *et al*., 2017; Farhat *et al*., 2019; Payne *et al*., 2019) provides strongest proof of the capacity for genetic variation in *M. tuberculosis*, other lines of evidence include the highly subdivided population structure of the *M. tuberculosis* Complex (Riojas *et al*., 2018), the well-described geographical host-pathogen sympatry (Hershberg *et al*., 2008; Brynildsrud *et al*., 2018) and, more recently, the observation of intra-patient bacillary microdiversity (Ley *et al*., 2019). In combination, these elements support the ongoing evolution of *M. tuberculosis*, as well as suggest the potential that “anti-evolution” therapeutics might yield much greater benefit in the clinical context than can be inferred from *in vitro* studies – in which the pressures on an obligate pathogen can only be approximated. That is, in addition to identifying the mutasome as target for adjunctive therapeutics designed to protect anti-TB drugs against emergent resistance, the results presented here support the further exploration of this and related strategies to disarm host-adaptive mechanisms in a major human pathogen.

## MATERIALS AND METHODS

### Bacterial strains and culture conditions

All mycobacterial strains (**Supplementary File 2 – Key Reagents**) were grown in liquid culture containing Difco™ Middlebrook 7H9 Broth (BD Biosciences, San Jose, CA) and supplemented with 0.2 % (v/v) glycerol (Sigma Aldrich, St. Louis, MO), 0.005 % (v/v) Tween® 80 (Sigma Aldrich, St. Louis, MO), and 10 % (v/v) BBL™ Middlebrook OADC Enrichment (BD Biosciences, San Jose, CA). For *M. smegmatis*, liquid cultures were incubated at 37 °C with orbital shaking at 100 rpm, until the desired growth density was attained – measured by spectrophotometry at a wavelength of 600 nm – before further experimentation. Solid media comprised Difco™ Middlebrook 7H10 Agar (BD Biosciences, San Jose, CA) supplemented with 0.5 % (v/v) glycerol (Sigma Aldrich, St. Louis, MO), and 10 % (v/v) BBL™ Middlebrook OADC Enrichment (BD Biosciences, San Jose, CA). Solid media plates were incubated at 37 °C for 3–4 days or until colonies had formed.

### Mutasome reporter constructs

The V-*imuA′* construct was designed by altering the coding sequence of *imuA′* within the complementing vector, pAINT::*imuA′* (Warner *et al*., 2010), so that the coding sequence of Venus fluorescent protein (VFP) (Nagai *et al*., 2002) was inserted in-frame after the start codon of the *imuA′* ORF. Furthermore, an in-frame FLAG tag sequence (Einhauer and Jungbauer, 2001) was inserted between the coding region of *vfp* and *imuA′* to produce a single ORF encoding VFP-FLAG-ImuA′. For ImuB, the construct PSOS(*imuA′*)-*egfp*-*imuB* was designed such that the regulatory elements immediately upstream of *imuA′* were inserted immediately upstream of the *imuB* ORF which was further altered by inserting the sequence encoding EGFP (Cormack *et al*., 1996) linked to a FLAG tag-encoded sequence immediately after the start codon of *imuB* to produce a single ORF encoding EGFP-FLAG-ImuB’ which was cloned into pMCAINT::*imuB* (Warner *et al*., 2010). The photoconvertible PSOS(*imuA′*)-*mEos4A-imuB* construct was based on the G-*imuB* construct, such that the coding sequence of EGFP was replaced by mEos4a (Paez Segala *et al*., 2015), yielding mEos4A-FLAG-ImuB. For DnaE2, the *egfp* sequence was inserted in-frame after the start codon of *M. smegmatis dnaE2*.

### Mutant binding G-*imuB*^AAAAG^ construct

To introduce the ^352^AAAAGG^357^ ImuB allele (Warner *et al*., 2010) into the EGFP-ImuB protein, the nucleotide sequence from pMCAINT::*imuB*^AAAAGG^ was swapped into the corresponding position of PSOS(*imuA′*)-*egfp-imuB* to yield pMCAINT::PSOS(*imuA′*)-*egfp-imuB*^AAAAGG^.

### *M. smegmatis* mutasome reporter strains

*M. smegmatis* strain V-*imuA′* was generated by introducing the pAINT::*vfp*-*imuA′* plasmid into Δ*imuA′* (Warner *et al*., 2010) by the standard electroporation method. Strains G-*imuB*, PC-*imuB*, and G-*imuB*^AAAAGG^ were developed by integration of the pMCAINT::PSOS(*imuA′*)-*egfp*-*imuB*, pMCAINT::PSOS(*imuA′*)-*mEos4a-imuB*, or pMCAINT::PSOS(*imuA′*)-*egfp*-*imuB*^AAAAGG^ plasmid, respectively, into the genome of Δ*imuB* (Warner *et al*., 2010). The *dnaN-mCherry*::G-*imuB* strain was developed by the electroporation of pMCAINT::PSOS(*imuA′*)-*egfp*-*imuB* into the *M. smegmatis dnaN-mCherry* background (Santi *et al*., 2013). To generate the G-dnaE2 strain, pTweety::*egfp-dnaE2* was electroporated into Δ*dnaE2* (Warner *et al*., 2010). Similarly, *dnaN-mCherry*::G-*dnaE2* was generated by electroporation of pTweety::*egfp-dnaE2* into *M. smegmatis dnaN-mCherry*. Mutasome-deficient strains Δ*imuA′*, Δ*dnaE2, and dnaE2^AIA^* were electroporated with pMCAINT::PSOS(*imuA′*)-*egfp*-*imuB* to produce Δ*imuA′*::*G-imuB*, Δ*dnaE2*::*G-imuB*, and *dnaE2*^AIA^::*G-imuB*, respectively.

### Mutagenesis Assays

Mutagenesis assays were performed as previously described (Boshoff *et al*., 2003; Warner *et al*.,2010), with RIF-resistant colonies enumerated on solid media after 5 days of growth. Mutation frequencies were calculated by dividing the number of RIF-resistant colonies of each sample by the CFU/ml of un-irradiated sample.

### Antibiotic treatments

MMC (Mitomycin C from *Streptomyces caespitosus*) (Sigma Aldrich, St. Louis, MO) was dissolved in ddH_2_O, while GRS was dissolved in DMSO. Cultures of *M. smegmatis* were grown in 7H9-OADC— supplemented with selection antibiotic where applicable—at 37 °C to an optical density (OD_600_) of between 0.2 and 0.4. Thereafter, cultures were split into different 5 ml cultures and MMC and/or GRS was added to a final concentration dependent on the MIC (Kling *et al*., 2015).

### Snapshot microscopy

Single snapshot micrographs of *M. smegmatis* cells were captured with a Zeiss Axioskop M, Zeiss Axio.Scope, and Zeiss Axio.Observer Z1. Briefly, 2.0–5.0 μl of liquid culture was placed between a No. 1.5 glass coverslip and microscope slide. A transmitted mercury lamp light was used together with filter cubes to visualize fluorescence using a 100× 1.4 NA plan apochromatic oil immersion objective lens. Samples were located using either transmitted light, differential interference contrast (DIC), or epifluorescence. Snapshot images were captured with either a Zeiss 1 MP or Zeiss AxioCam HRm monochrome camera. Images of the same experiment were captured with the same instrument and exposure settings. Green fluorescence of EGFP was detected using the Zeiss Filter Set 38 HE. Red fluorescence of mCherry was detected using the Zeiss Filter Set 43. Images were captured using AxioVision 4.7 or ZEN Blue Microscope and Imaging Software. Images were processed using Fiji (Schindelin *et al*., 2012); images of the same strain were contrasted to the same maximum and minimum within an experiment.

### Quantitative image analysis

*M. smegmatis* bacilli were plotted from shortest to longest and aligned according to their midcell position (0 on the y axis) using the MicrobeJ plugin of ImageJ (Ducret *et al*., 2016). Along each point of the cell, a dot was generated and coloured according to the fluorescence intensity along the medial axis of the bacillus. Therefore, this plot represents the fluorescence intensity along the medial axis of every bacillus imaged under the relevant experimental conditions. R was used for visual representation of the data.

### Super-resolution iPALM microscopy

Three-dimensional PALM was performed using the iPALM instrument (Shtengel *et al*., 2009). Round 25 mm diameter No. 1.5 coverslips were cleaned by sonication in 1 M KOH for 45 minutes. Following rinsing in deionized water and drying at 60 °C, coverslips were coated with 5mg/ml >70,000 molecular weight poly-L-lysine hydrobromide (MP Biomedicals, Santa Ana, CA) for 30 minutes at room temperature. Thereafter, gold nanorods were adhered to the coverslips for 30 minutes before drying by vacuum centrifugation. Thin film deposition was used to coat the fiducial coverslips with SiO_2_. Thereafter, the fiducial coverslips were cleaned with 1 M KOH and coated with 1% poly-L-lysine for 60 minutes at 37.0 °C. Bacterial cultures of OD_600_ = 0.4 were exposed to drug conditions (5× MMC or GRS) for 6 h before 3.0 ml of bacterial sample was centrifuged onto a fiducial coverslip at 3,200 rcf for 15 minutes in a six-well plate. The sample was rinsed three times in Dulbecco’s PBS and fixed with 0.5% paraformaldehyde for 2 minutes. Thereafter, the sample was mounted in Dulbecco’s PBS and the gold coverslip were adhered to a clean (as above) 18mm diameter No. 1.5 round coverslip. Each coverslip was sealed to prevent evaporation. The sample was mounted between two opposing Nikon 60× 1.49 NA Apo TIRF oil immersion lenses and captured using three Andor iXon-3 EMCCD cameras. Bacterial cells were located by DIC visualization, and each sample was imaged three times at separate regions containing 2–5 bacilli using TIRF illumination. Experiments were repeated at least three times. A calibration image of 100 cycles was taken of the gold fiducials in each field-of-view. Bacilli were imaged for 25,000 cycles using alternating 405 nm (mEos4a photoconversion) and 561 nm (converted mEos4a excitation) laser cycles per frame. FF01-593/40 emission filters (Semrock) were used during mEos4a imaging. Thereafter, the calibration file was processed using PeakSelector™ (Shtengel *et al*., 2009) and used to calibrate the detected localizations. During image processing, low confidence localizations were excluded based on unwrapped Z-error and Z-position. Images were produced with PeakSelector™.

### Single-cell time-lapse fluorescence microscopy

Liquid cultures of *M. smegmatis* reporter strains were grown to mid-logarithmic phase (OD_600_ = 0.6), cells were collected by centrifugation at 3900 × *g* for 5 min and concentrated 10-fold in 7H9 medium. The cells were filtered through a polyvinylidene difluoride (PVDF) syringe filter (Millipore) with a 5 μm pore size to yield a clump-free cell suspension. The single cell suspension was spread on a semi-permeable membrane and secured between a glass coverslip and the *serpentine 2 chip* (Delincé *et al*.,2016) in a custom-made PMMA/Aluminium holder (Dhar and Manina, 2015). Time-lapse microscopy employing a DeltaVision personalDV inverted fluorescence microscope (Applied Precision, WA) with a 100x oil immersion objective was used to image single cells of *M. smegmatis*. The bacteria and microfluidic chip were maintained at 37 °C in an environmental chamber with a continuous flow of 7H9 medium, with or without 100 ng/ml of MMC, at a constant flow rate of 25 μl.min^-1^, as described previously (Wakamoto *et al*., 2013; Dhar and Manina, 2015). Images were obtained every 10 min on phase-contrast and fluorescence channels (for EGFP, excitation filter 470/40 nm, emission filter 525/50 nm; for mCherry, excitation filter 572/35, emission filter 632/60; for YFP excitation filter 500/20 nm, emission filter 535/30 nm) using a CoolSnap HQ2 camera. Image-based autofocus was performed on each point prior to image acquisition. Experiments were repeated 2–4 times; a typical experiment collected images from up to 80 XY points at the 10 min intervals. The images were analysed using Fiji (Schindelin *et al*., 2012).

### Protein expression and purification

N-terminally His-tagged *M. smegmatis* ImuB was co-expressed with ImuA′ in *E. coli* BL21(DE3) cells using two expression vectors from the NKI-LIC vector suite (Luna-Vargas *et al*., 2011): pETNKI-his-3C-LIC-kan for ImuB and pCDFNKI-StrepII3C-LIC-strep for ImuA’ that have different resistance markers, kanamycin and streptomycin; as well as different origins of replication, ColE1 and CloDF13, respectively. Protein production was induced with isopropyl 1-thio-β-d-galactopyranoside (IPTG) at 30°C for 2 hours. The ImuBA′ complex was purified using a Histrap column followed by a Superdex 200 16/60 column. Both N-His6 *M. smegmatis* ImuB and β clamp were expressed in *E. coli* BL21(DE3) cells and purified using HisTrap, HiTrap Q, and S200 columns. All proteins were flash frozen in liquid nitrogen and stored at −80 °C.

### Size-exclusion chromatography analysis

Samples of individual proteins and the different complexes were injected onto a PC3.2/30 (2.4 ml) Superdex 200 Increase gel filtration column (GE Healthcare) pre-equilibrated in 50 mM Tris pH 8.5 and 300 mM NaCl. Thereafter, 50 μl fractions were collected and analyzed by SDS–PAGE electrophoresis using 4–12% NuPage Bis-Tris precast gels (Life Technologies). Gels were stained with 0.01% (v/v) 2,2,2-Trichloroethanol (TCE) and imaged with UV light.

### Thermal unfolding experiments

Melting curves of the *M. smegmatis* β clamp (5μM) in the presence and absence of GRS (15μM) were measured in UV-capillaries using the Tycho NT6 (NanoTemper Technologies) where the protein unfolding is followed by detecting the fluorescence of intrinsic tryptophan and tyrosine residues at both emission wavelengths of 350 nm and 330 nm.

## ACKNOWLEDGMENTS

This work was supported by the US National Institute of Child Health and Human Development (NICHD) U01HD085531 (to D.F.W. and R.W.). We acknowledge the funding support of the Research Council of Norway (R&D Project 261669 “Reversing antimicrobial resistance”) (to D.F.W.), the South African Medical Research Council (to V.M. and D.F.W.); the National Research Foundation of South Africa (to D.F.W. and V.M.); a Senior International Research Scholars grant from the Howard Hughes Medical Institute (to V.M.); and a LUMC Fellowship (to M.H.L.). In addition, M.A.R. is grateful to the South African National Research Foundation (NRF) for financial assistance during his PhD training (grant no. 104683) as well as the Whitehead Scientific Travel Award. Z.A.M is grateful to the University of Cape Town, the David and Elaine Potter Foundation Research Fellowship, and the Swiss Government Excellence Research Scholarship for financial assistance during her PhD.

## SUPPLEMENTARY DATA

**Supplementary File 1 – Supplementary figures**

**Supplementary File 2 – Key reagents**

**Video 1 – Time-lapse microscopy of G-ImuB and mCherry-DnaN dual reporter.** Representative time-lapse movie of the reporter strain expressing G-ImuB and mCherry-DnaN. Bacteria were imaged on fluorescence and phase channels for up to 36 hours at 10-minute intervals. Treatment with MMC (100 ng/ml) was at 0 – 4.5 hours. This experiment was repeated 6 times. Numbers indicate the hours (h) elapsed in the time-lapse experiment. 7H9, Middlebrook 7H9/OADC. MMC, Mitomycin C.Scale bar, 5 μm. G-ImuB, green; mCherry-DnaN, magenta; overlay, white.

**Video 2 – Time-lapse microscopy of V-ImuA**′ **and mCherry-DnaN dual reporter.** Representative time-lapse movie of the reporter strain expressing V-ImuA′ and mCherry-DnaN. Bacteria were imaged on fluorescence and phase channels for up to 36 hours at 10-minute intervals. Treatment with MMC (100 ng/ml) was at 0 – 4.5 hours. This experiment was repeated 3 times. Numbers indicate the hours (h) elapsed in the time-lapse experiment. 7H9, Middlebrook 7H9/OADC. MMC, Mitomycin C. Scale bar, 5 μm. V-ImuA′, green; mCherry-DnaN, magenta; overlay, white.

**Video 3 – Time-lapse microscopy of G-DnaE2 and mCherry-DnaN dual reporter.** Representative time-lapse movie of the reporter strain expressing G-DnaE2 and mCherry-DnaN. Bacteria were imaged on fluorescence and phase channels for up to 36 hours at 10-minute intervals. Treatment with MMC (100 ng/ml) was at 0 – 4.5 hours. This experiment was repeated 3 times. Numbers indicate the hours (h) elapsed in the time-lapse experiment. 7H9, Middlebrook 7H9/OADC. MMC, Mitomycin C. Scale bar, 5 μm. G-DnaE2, green; mCherry-DnaN, magenta; overlay, white.

## REFERENCES

Adefisayo, O.O., Dupuy, P., Bean, J.M. and Glickman, M.S. (2021). Division of labor between SOS and PafBC in mycobacterial DNA repair and mutagenesis. bioRxiv, p. 2021.08.05.455301. doi:10.1101/2021.08.05.455301.

Bargonetti, J., Champeil, E. and Tomasz, M. (2010). Differential toxicity of DNA adducts of mitomycin C. Journal of Nucleic Acids, 698960. doi:10.4061/2010/698960.

Barrett, T.C., Mok, W.W.K.K., Murawski, A.M. and Brynildsen, M.P. (2019). Enhanced antibiotic resistance development from fluoroquinolone persisters after a single exposure to antibiotic. Nature Communications, 10(1), pp. 1–11. doi:10.1038/s41467-019-09058-4.

Boritsch, E.C. and Brosch, R. (2016). Evolution of *Mycobacterium tuberculosis*: New insights into pathogenicity and drug resistance. Microbiology Spectrum, (5), pp. 495–515. doi:10.1128/9781555819569.ch22.

Bosch, B., DeJesus, M.A., Poulton, N.C., Zhang, W., Engelhart, C.A., Zaveri, A., Lavalette, S., Ruecker, N., Trujillo, C., Wallach, J.B., Li, S., Ehrt, S., Chait, B.T., Schnappinger, D. and Rock, J.M. (2021). Genome-wide gene expression tuning reveals diverse vulnerabilities of *M. tuberculosis*. Cell, 184(17), pp. 4579–4592.e24. doi:10.1016/J.CELL.2021.06.033.

Boshoff, H.I. and Mizrahi, V. (2000). Expression of *Mycobacterium smegmatis* pyrazinamidase in *Mycobacterium tuberculosis* confers hypersensitivity to pyrazinamide and related amides. Journal of Bacteriology, 182(19), pp. 5479–85. doi:10.1128/JB.182.19.5479-5485.2000.

Boshoff, H.I.M.M., Reed, M.B., Barry, C.E. and Mizrahi, V. (2003). DnaE2 polymerase contributes to *in vivo* survival and the emergence of drug resistance in *Mycobacterium tuberculosis*. Cell, 113(2), pp. 183–193. doi:10.1016/S0092-8674(03)00270-8.

Brynildsrud, O.B., Pepperrell, C.S., Suffys, P., Grandjean, L., Monteserin, J., Debech, N., Bohlin, J., Alfsnes, K., Pettersson, J.O.H., Kirkeleite, I., Fandinho, F., da Silva, M.A., Perdigao, J., Portugal, I., Viveiros, M., Clark, T., Caws, M., Dunstan, S., Thai, P.V.K., Lopez, B., Ritacco, V., Kitchen, A., Brown, T.S., van Soolingen, D., O’Neill, M.B., Holt, K.E., Feil, E.J., Mathema, B., Balloux, F., Eldholm, V. (2018). Global expansion of *Mycobacterium tuberculosis* lineage 4 shaped by colonial migration and local adaptation. Science Advances, 4(10), pp. 1–12. doi:10.1126/sciadv.aat5869.

Bunting, K.A., Roe, S.M. and Pearl, L.H. (2003). Structural basis for recruitment of translesion DNA polymerase Pol IV/DinB to the β-clamp. EMBO Journal, 22(21), pp. 5883–5892. doi:10.1093/emboj/cdg568.

Burnouf, D.Y., Olieric, V., Wagner, J., Fujii, S., Reinbolt, J., Fuchs, R.P.P. and Dumas, P. (2004). Structural and biochemical analysis of sliding clamp/ligand interactions suggest a competition between replicative and translesion DNA polymerases. Journal of Molecular Biology, 335(5), pp. 1187–1197. doi:10.1016/j.jmb.2003.11.049.

Chengalroyen, M.D., Mason, M.K., Borsellini, A., Tassoni, R., Abrahams, G.L., Lynch, S., Ahn, Y.-M., Ambler, J., Young, K., Crowley, B.M., Olsen, D.B., Warner, D.F., Barry, C.E., Boshoff, H.I.M., Lamers, M.H. and Mizrahi, V. (2021) DNA-dependent binding of nargenicin to DnaE1 inhibits replication in *Mycobacterium tuberculosis*. bioRxiv, p. 2021.10.27.466036. doi:10.1101/2021.10.27.466036.

Cirz, R.T., Chin, J.K., Andes, D.R., Crécy-Lagard, V. de, Craig, W.A. and Romesberg, F.E. (2005). Inhibition of mutation and combating the evolution of antibiotic resistance. PLOS Biology, 3(6), p. e176. doi:10.1371/JOURNAL.PBIO.0030176.

Cokol, M., Kuru, N., Bicak, E., Larkins-Ford, J. and Aldridge, B.B. (2017). Efficient measurement and factorization of high-order drug interactions in *Mycobacterium tuberculosis*. Science Advances, 3(10). doi:10.1126/sciadv.1701881.

Cormack, B.P., Valdivia, R.H. and Falkow, S. (1996). FACS-optimized mutants of the green fluorescent protein (GFP). Gene, 173(1), pp. 33–38. doi:10.1016/0378-1119(95)00685-0.

Delincé, M.J., Bureau, J.-B., López-Jiménez, A.T., Cosson, P., Soldati, T. and McKinney, J.D. (2016). A microfluidic cell-trapping device for single-cell tracking of host–microbe interactions. Lab Chip, 16(17), pp. 3276–3285. doi:10.1039/C6LC00649C.

Dhar, N. and Manina, G. (2015). Single-cell analysis of mycobacteria using microfluidics and time-lapse microscopy. Methods in Molecular Biology, 1285, pp. 241–256.

Ducret, A., Quardokus, E. and Brun, Y. (2016). MicrobeJ, a tool for high throughput bacterial cell detection and quantitative analysis. Nature Microbiology, 1(7), pp. 1–14. doi:10.1038/nmicrobiol.2016.77.MicrobeJ.

van Eijk, E., Wittekoek, B., Kuijper, E.J. and Smits, W.K. (2017). DNA replication proteins as potential targets for antimicrobials in drug-resistant bacterial pathogens. Journal of Antimicrobial Chemotherapy, 72(5), pp. 1275–1284. doi:10.1093/JAC/DKW548.

Einhauer, A. and Jungbauer, A. (2001). The FLAG™ peptide, a versatile fusion tag for the purification of recombinant proteins. Journal of Biochemical and Biophysical Methods, 49(1–3), pp. 455–465. doi:10.1016/S0165-022X(01)00213-5.

Erdem, A.L., Jaszczur, M., Bertram, J.G., Woodgate, R., Cox, M.M. and Goodman, M.F. (2014). DNA polymerase V activity is autoregulated by a novel intrinsic DNA-dependent ATPase. eLife, 2014(3). doi:10.7554/ELIFE.02384.

Erill, I., Campoy, S., Mazon, G. and Barbé, J. (2006). Dispersal and regulation of an adaptive mutagenesis cassette in the bacteria domain. Nucleic Acids Research, 34(1), pp. 66–77. doi:10.1093/nar/gkj412.

Farhat, M.R., Freschi, L., Calderon, R., Ioerger, T., Snyder, M., Meehan, C.J., de Jong, B., Rigouts, L., Sloutsky, A., Kaur, D., Sunyaev, S., van Soolingen, D., Shendure, J., Sacchettini, J. and Murray, M. (2019). GWAS for quantitative resistance phenotypes in *Mycobacterium tuberculosis* reveals resistance genes and regulatory regions. Nature Communications, 10(1). doi:10.1038/s41467-019-10110-6.

Franklin, W.A., Doetsch, P.W. and Haseltine, W.A. (1985). Structural determinatlon of the ultraviolet light-induced thymine-cytosine pyrimidine-pyrimidone (6–4) photoproduct. Nucleic Acids Research, 13(14), pp. 5317–5325. doi:10.1093/NAR/13.14.5317.

Gagneux, S. (2018). Ecology and evolution of *Mycobacterium tuberculosis*. Nature Reviews Microbiology, 16(4), pp. 202–213. doi:10.1038/nrmicro.2018.8.

Galagan, J.E. (2014). Genomic insights into tuberculosis. Nature Reviews Genetics, 15(5), pp. 307–320. doi:10.1038/nrg3664.

Goodman, M.F., McDonald, J.P., Jaszczur, M.M. and Woodgate, R. (2016) Insights into the complex levels of regulation imposed on *Escherichia coli* DNA polymerase V. DNA Repair, 44, pp. 42–50. doi:10.1016/J.DNAREP.2016.05.005.

Gray, T.A. and Derbyshire, K.M. (2018). Blending genomes: distributive conjugal transfer in mycobacteria, a sexier form of HGT. Molecular Microbiology, 108(6), pp. 601–613. doi:10.1111/mmi.13971.

Gruber, A.J., Erdem, A.L., Sabat, G., Karata, K., Jaszczur, M.M., Vo, D.D., Olsen, T.M., Woodgate, R., Goodman, M.F. and Cox, M.M. (2015). A RecA protein surface required for activation of DNA polymerase V. PLOS Genetics, 11(3), p. e1005066. doi:10.1371/JOURNAL.PGEN.1005066.

Hershberg, R., Lipatov, M., Small, P.M., Sheffer, H., Niemann, S., Homolka, S., Roach, J.C., Kremer, K., Petrov, D.A., Feldman, M.W. and Gagneux, S. (2008). High functional diversity in *Mycobacterium tuberculosis* driven by genetic drift and human demography. PLoS Biology, 6(12), pp. 2658–2671. doi:10.1371/journal.pbio.0060311.

Ippoliti, P.J., DeLateur, N.A., Jones, K.M. and Beuning, P.J. (2012). Multiple strategies for translesion synthesis in bacteria. Cells, 1(4), pp. 799–831. doi:10.3390/cells1040799.

Jaszczur, M.M., Vo, D.D., Stanciauskas, R., Bertram, J.G., Sikand, A., Cox, M.M., Woodgate, R., Mak, C.H., Pinaud, F. and Goodman, M.F. (2019). Conformational regulation of *Escherichia coli* DNA polymerase V by RecA and ATP. PLoS Genetics, 15(2), pp. 1–27. doi:10.1371/journal.pgen.1007956.

Jiang, Q., Karata, K., Woodgate, R., Cox, M.M. and Goodman, M.F. (2009). The active form of DNA polymerase V is UmuD′2C-RecA-ATP. Nature, 460(7253), pp. 359–363. doi:10.1038/nature08178.

Joseph, A.M., Daw, S., Sadhir, I. and Badrinarayanan, A. (2021). Coordination between nucleotide excision repair and specialized polymerase Dnae2 action enables dna damage survival in non-replicating bacteria. eLife, 10. doi:10.7554/ELIFE.67552.

Kling, A., Lukat, P., Almeida, D.V., Bauer, A., Fontaine, E., Sordello, S., Zaburannyi, N., Herrmann, J., Wenzel, S., Konig, C., Ammerman, N.C., Barrio, M.B., Borchers, K., Bordon-Pallier, F., Bronstrup, M., Courtemanche, G., Gerlitz, M., Geslin, M., Hammann, P., Heinz, D.W., Hoffmann, H., Klieber, S., Kohlmann, M., Kurz, M., Lair, C., Matter, H., Nuermberger, E., Tyagi, S., Fraisse, L., Grosset, J., H., Lagrange, S., Müller, R., (2015). Targeting DnaN for tuberculosis therapy using novel griselimycins. Science, 348(6239), pp. 1106–1112. doi:10.1126/science.aaa4690.

Ley, S.., de Vos, M., Van Rie, A. and Warren, R.M. (2019.) Deciphering within-host microevolution of *Mycobacterium tuberculosis* through whole-genome sequencing: the phenotypic impact and way forward. Microbiology and Molecular Biology Reviews, 83(2), pp. 1–21.

Lin, P.L. and Flynn, J.L. (2018). The end of the binary era: Revisiting the spectrum of tuberculosis. The Journal of Immunology, 201(9), pp. 2541–2548. doi:10.4049/JIMMUNOL.1800993.

Luna-Vargas, M.P.A., Christodoulou, E., Alfieri, A., van Dijk, W.J., Stadnik, M., Hibbert, R.G., Sahtoe, D.D., Clerici, M., Marco, V. De, Littler, D., Celie, P.H.N., Sixma, T.K. and Perrakis, A. (2011) Enabling high-throughput ligation-independent cloning and protein expression for the family of ubiquitin specific proteases. Journal of Structural Biology, 175(2), pp. 113–119. doi:10.1016/J.JSB.2011.03.017.

Maslowska, K.H., Makiela-Dzbenska, K. and Fijalkowska, I.J. (2019) The SOS system: A complex and tightly regulated response to DNA damage. Environmental and Molecular Mutagenesis, 60(4), pp. 368–384. doi:10.1002/em.22267.

McHenry, C.S. (2011). Breaking the rules: bacteria that use several DNA polymerase IIIs. EMBO reports, 12(5), pp. 408–414. doi:10.1038/embor.2011.51.

Merrikh, H. and Kohli, R.M. (2020). Targeting evolution to inhibit antibiotic resistance. FEBS Journal, 287(20), pp. 4341–4353. doi:10.1111/febs.15370.

Minias, A., Brzostek, A. and Dziadek, J.J. (2018). Targeting DNA repair systems in antitubercular drug development. Current Medicinal Chemistry, 25(8), pp. 1–12. doi:10.2174/0929867325666180129093546.

Mittal, P., Sinha, R., Kumar, A., Singh, P., Ngasainao, M.R., Singh, A. and Singh, I.K. (2020). Focusing on DNA repair and damage tolerance mechanisms in *Mycobacterium tuberculosis*: An emerging therapeutic theme. Current Topics in Medicinal Chemistry, 20(5), pp. 390–408. doi:10.2174/1568026620666200110114322.

Nagai, T., Ibata, K., Park, E.S., Kubota, M., Mikoshiba, K. and Miyawaki, A. (2002). A variant of yellow fluorescent protein with fast and efficient maturation for cell-biological applications. Nature Biotechnology, 20(1), pp. 87–90. doi:10.1038/nbt0102-87.

Paez Segala, M.G., Sun, M., Shtengel, G., Viswanathan, S., Baird, M., Macklin, J., Patel, R., Allen, J., Howe, E., Piszczek, G., Hess, H., Davidson, M., Wang, Y. and Looger, L. (2015). Fixation-resistant photoactivatable fluorescent proteins for correlative light and electron microscopy. Nature methods, 12(3), pp. 215–218. doi:10.1038/nmeth.3225.

Payne, J.L., Menardo, F., Trauner, A., Borrell, S., Gygli, S.M., Loiseau, C., Gagneux, S. and Hall, A.R. (2019). Transition bias influences the evolution of antibiotic resistance in *Mycobacterium tuberculosis*. PLoS, 17(5), pp. 1–23. doi:10.1101/421651.

Ragheb, M.N., Thomason, M.K., Hsu, C., Nugent, P., Gage, J., Samadpour, A.N., Kariisa, A., Merrikh, C.N., Miller, S.I., Sherman, D.R. and Merrikh, H. (2019). Inhibiting the evolution of antibiotic resistance. Molecular Cell, 73(1), pp. 157–165.e5. doi:10.1016/j.molcel.2018.10.015.

Reiche, M.A., Warner, D.F. and Mizrahi, V. (2017). Targeting DNA replication and repair for the development of novel therapeutics against tuberculosis. Frontiers in Molecular Biosciences, 4(November), pp. 1–18. doi:10.3389/fmolb.2017.00075.

Renzette, N., Gumlaw, N., Nordman, J.T., Krieger, M., Yeh, S.P., Long, E., Centore, R., Boonsombat, R. and Sandler, S.J. (2005). Localization of RecA in *Escherichia coli* K-12 using RecA-GFP. Molecular Microbiology, 57(4), pp. 1074–1085. doi:10.1111/j.1365-2958.2005.04755.x.

Revitt-Mills, S.A. and Robinson, A. (2020). Antibiotic-induced mutagenesis: Under the microscope. Frontiers in Microbiology, 0, p. 2611. doi:10.3389/FMICB.2020.585175.

Riojas, M.A., McGough, K.J., Rider-Riojas, C.J., Rastogi, N. and Hazbón, M.H. (2018). Phylogenomic analysis of the species of the *Mycobacterium tuberculosis* complex demonstrates that *Mycobacterium africanum*, *Mycobacterium bovis*, *Mycobacterium caprae*, *Mycobacterium microti* and *Mycobacterium pinnipedii* are later heterotypic synonyms of *Mycobacterium tuberculosis*. International Journal of Systematic and Evolutionary Microbiology, 68(1), pp. 324–332. doi:10.1099/IJSEM.0.002507.

Robinson, A., McDonald, J.P., Caldas, V.E.A., Patel, M., Wood, E.A., Punter, C.M., Ghodke, H., Cox, M.M., Woodgate, R., Goodman, M.F., van Oijen, A.M. and Oijen, A.M. (2015). Regulation of mutagenic DNA polymerase V activation in space and time. PLOS Genetics, 11(8), p. e1005482.

Santi, I., Dhar, N., Bousbaine, D., Wakamoto, Y. and McKinney, J.D. (2013). Single-cell dynamics of the chromosome replication and cell division cycles in mycobacteria. Nature Communications, 4(May), pp. 1–10. doi:10.1038/ncomms3470.

Santi, I. and McKinney, J.D. (2015). Chromosome organization and replisome dynamics in *Mycobacterium smegmatis*. mBio, 6(1). doi:10.1128/mBio.01999-14.

Schindelin, J., Arganda-Carreras, I., Frise, E., Kaynig, V., Longair, M., Pietzsch, T., Preibisch, S., Rueden, C., Saalfeld, S., Schmid, B., Tinevez, J.-Y., White, D.J., Hartenstein, V., Eliceiri, K., Tomancak, P. and Cardona, A. (2012). Fiji: an open-source platform for biological-image analysis. Nature Methods, 9(7), pp. 676–682. doi:10.1038/nmeth.2019.

Shtengel, G., Galbraith, J.A., Galbraith, C.G., Lippincott-Schwartz, J., Gillette, J.M., Manley, S., Sougrat, R., Waterman, C.M., Kanchanawong, P., Davidson, M.W., Fetter, R.D. and Hess, H.F. (2009). Interferometric fluorescent super-resolution microscopy resolves 3D cellular ultrastructure. Proceedings of the National Academy of Sciences of the United States of America, 106(9), pp. 3125–3130. doi:10.1073/pnas.0813131106.

Singh, A. (2017). Guardians of the mycobacterial genome: A review on DNA repair systems in *Mycobacterium tuberculosis*. Microbiology, 163(12), pp. 1740–1758. doi:10.1099/mic.0.000578.

Smith, P.A. and Romesberg, F.E. (2007). Combating bacteria and drug resistance by inhibiting mechanisms of persistence and adaptation. Nature Chemical Biology, 3(9), pp. 549–556. doi:10.1038/nchembio.2007.27.

Timinskas, K., Balvočiute, M., Timinskas, A. and Venclovas, Č. (2014). Comprehensive analysis of DNA polymerase III α subunits and their homologs in bacterial genomes. Nucleic Acids Research, 42(3), pp. 1393–1413. doi:10.1093/nar/gkt900.

Timinskas, K. and Venclovas, Č. (2019). New insights into the structures and interactions of bacterial Y-family DNA polymerases. Nucleic Acids Research, 47(9), pp. 4383–4405. doi:10.1093/nar/gkz198.

Tomasz, M. (1995). Mitomycin C: small, fast and deadly (but very selective). Chemistry and Biology, pp. 575–579. doi:10.1016/1074-5521(95)90120-5.

Veening, J.-W. and Blokesch, M. (2017). Interbacterial predation as a strategy for DNA acquisition in naturally competent bacteria. Nature Reviews Microbiology, 15(10), pp. 621–629. doi:10.1038/nrmicro.2017.66.

Wakamoto, Y., Dhar, N., Chait, R., Schneider, K., Signorino-Gelo, F., Leibler, S. and McKinney, J.D. (2013). Dynamic persistence of antibiotic-stressed mycobacteria. Science, 339, pp. 91–95.

Warner, D.F., Koch, A. and Mizrahi, V. (2015). Diversity and disease pathogenesis in *Mycobacterium tuberculosis*. Trends in Microbiology, pp. 14–21. doi:10.1016/j.tim.2014.10.005.

Warner, D.F., Ndwandwe, D.E., Abrahams, G.L., Kana, B.D., Machowski, E.E., Venclovas, Č. and Mizrahi, V. (2010). Essential roles for *imuA′-* and *imuB*-encoded accessory factors in DnaE2-dependent mutagenesis in *Mycobacterium tuberculosis*. Proceedings of the National Academy of Sciences of the United States of America, 107(29), pp. 13093–13098. doi:10.1073/pnas.1002614107.

Warner, D.F., Rock, J.M., Fortune, S.M. and Mizrahi, V. (2017). DNA replication fidelity in the *Mycobacterium tuberculosis* complex. In Gagneux, S. (eds) Strain variation in the Mycobacterium tuberculosis complex: Its role in biology, epidemiology and control. Advances in Experimental Medicine and Biology, 1019. Springer, Cham. doi:10.1007/978-3-319-64371-7.

World Health Organization (2021) Global Tuberculosis Report

von Wintersdorff, C.J.H., Penders, J., van Niekerk, J.M., Mills, N.D., Majumder, S., van Alphen, L.B., Savelkoul, P.H.M. and Wolffs, P.F.G. (2016). Dissemination of antimicrobial resistance in microbial ecosystems through horizontal gene transfer. Frontiers in Microbiology, 0(FEB), p. 173. doi:10.3389/FMICB.2016.00173.

