## Supplementary Figures for "The mycobacterial mutasome: composition and recruitment in live cells"

[1]SAMRC/NHLS/UCT Molecular Mycobacteriology Research Unit, DSI/NRF Centre of Excellence for Biomedical TB Research, Department of Pathology, University of Cape Town, South Africa; [2]Institute of Infectious Disease and Molecular Medicine, University of Cape Town, South Africa; [3]Laboratory of Microbiology and Microsystems, School of Life Sciences, Swiss Federal Institute of Technology in Lausanne (EPFL), Lausanne, Switzerland; [4]Advanced Imaging Center, Howard Hughes Medical Institute, United States of America; [5]Department of Cell and Chemical Biology, Leiden University Medical Center, The Netherlands; [6]Department of Integrative Biomedical Sciences, University of Cape Town, South Africa; [7]Confocal and Light Microscope Imaging Facility, Department of Human Biology, University of Cape Town, South Africa; [8]Helmholtz Institute for Pharmaceutical Research Saarland (HIPS), Helmholtz Centre for Infection Research, Germany; [9]German Centre for Infection Research (DZIF), Partner Site Hannover-Braunschweig, Germany; [10]Laboratory of Genomic Integrity, National Institute of Child Health and Human Development, National Institutes of Health, United States of America; [11]Wellcome Centre for Infectious Diseases Research in Africa, University of Cape Town, South Africa.

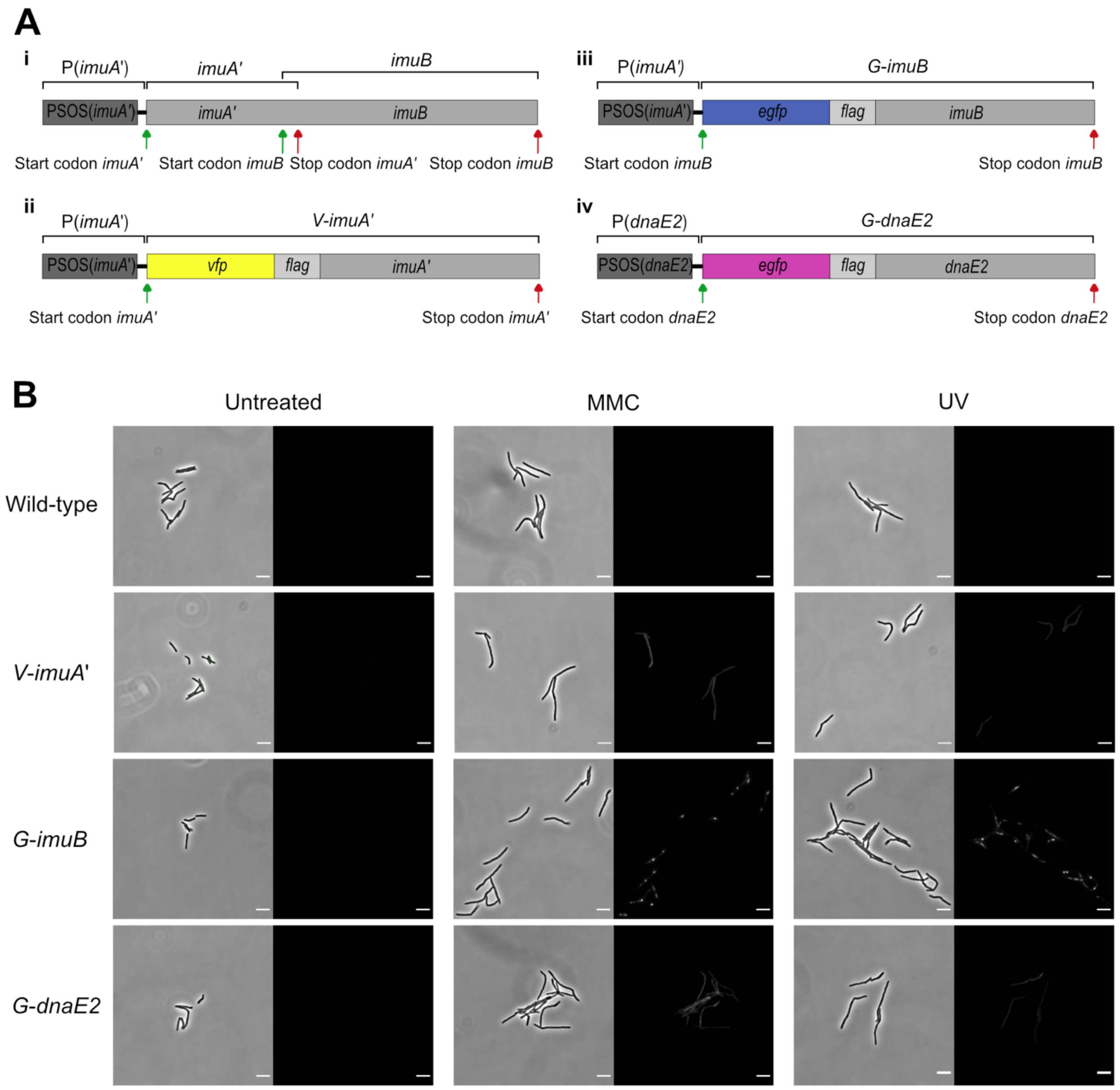


**Figure 1 – figure supplement 1. Design and representative fluorescent images of the mutasome translational reporters. A.** Construction of the reporter mutants, indicating: (i) the genomic structure of the wild-type *M. smegmatis imuA′-imuB* operon; (ii) the design of the *vfp-imuA′* gene utilizing the native *imuA′* promoter to control expression of a single VFP-ImuA′-encoding ORF; (iii) the PSOS*(imuA′)*-*egfp-imuB* construct encoding EGFP-ImuB; and (iv) the *egfp-dnaE2* allele used in the GFP-DnaE2 reporter. **B.** Representative brightfield and fluorescent images of wild-type, V-imuA′, G-imuB, and G-dnaE2 translational reporter strains untreated (left panel), following exposure to 1×MIC MMC (middle panel), or UV (right panel). Scale bars, 5 μm.

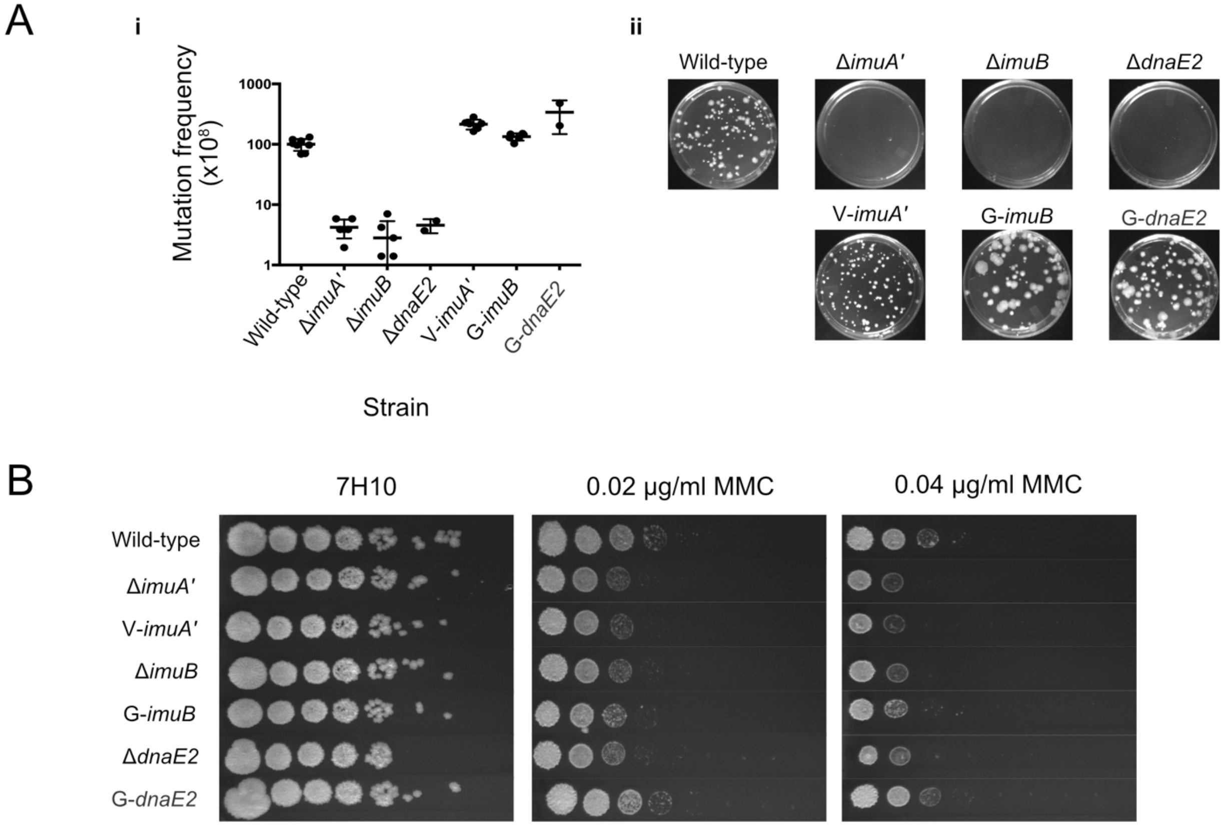


**Figure 1 – figure supplement 2. Functional validation of translational reporters. A.** N-terminally-tagged fluorescence reporter mutants of *M. smegmatis* ImuA′, ImuB, and DnaE2 retain function in DNA damage-induced mutagenesis. Cultures of *M. smegmatis* deletion mutants and complemented derivatives were exposed to 25 mJ/cm$^2$ of 254 nm UV light and allowed to recover for 3 hours before selection of rifampicin-resistant mutants on rifampicin-containing 7H10 solid agar plates. (i) Mutation frequencies were calculated as a fraction of the CFU/ml of each culture prior to exposure to UV irradiation. Complementation with the corresponding fluorescence reporter alleles restored the resistance frequencies of the three mutasome knockout mutants (Δ*imuA′*, Δ*imuB*, Δ*dnaE2*) to levels observed in wild-type *M. smegmatis*. (ii) Representative rifampicin-containing plates with rifampicin-resistant mutants. **B.** Serial dilutions of *M. smegmatis* deletion mutants and complemented strains were spotted on MMC containing plates. Results represent a minimum of three replicates for each strain.

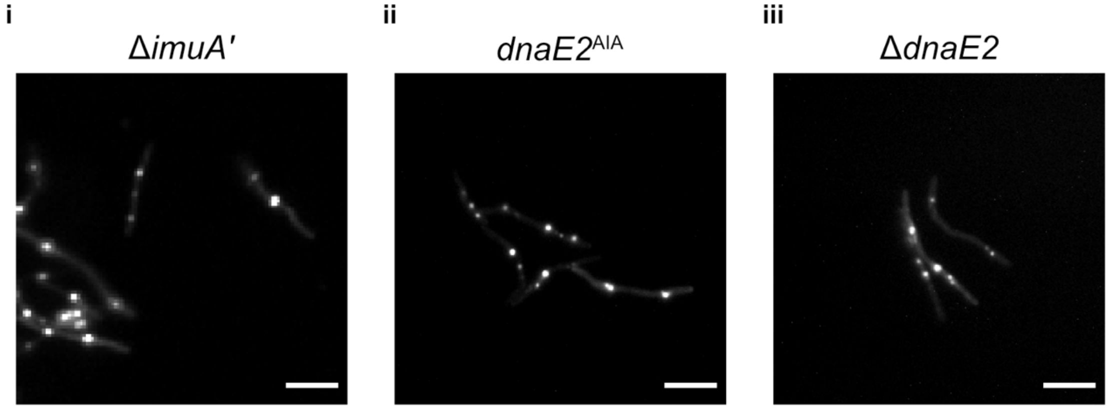


**Figure 2 – figure supplement 1. Formation of fluorescent EGFP-ImuB foci in the absence of functional mutasome components.** Each panel represents the expression of EGFP-ImuB foci following 1×MIC MMC treatment in different genetic backgrounds: (i) Δ*imuA′*, (ii) *dnaE2*$^{\mathrm{AIA}}$ and (iii) Δ*dnaE2*. Scale bars, 5 μm.

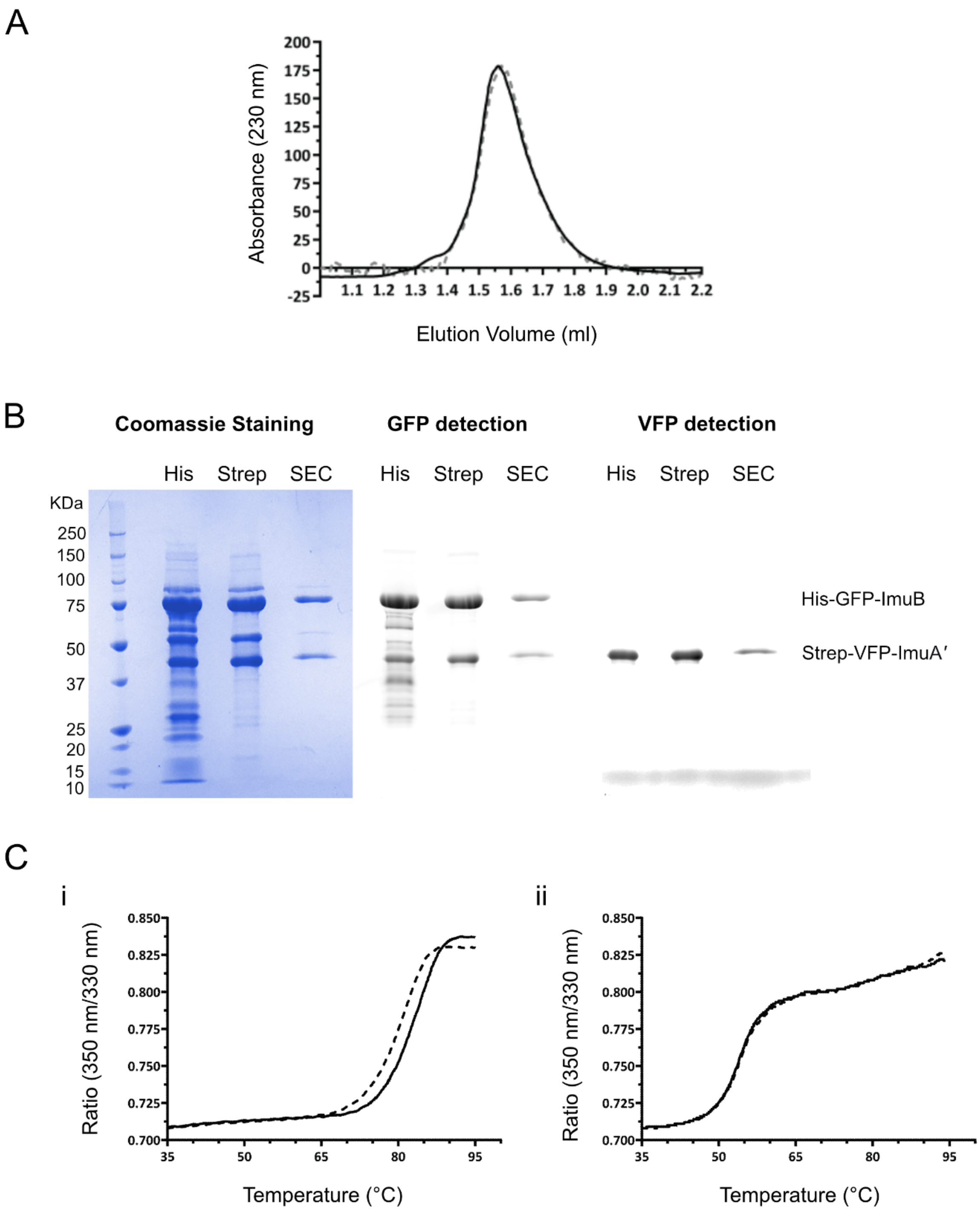


**Figure 3 – figure supplement 1. Biochemical confirmation of stable ImuA′-ImuB complex formation and GRS binding DnaN. A.** *M. smegmatis* ImuA′ and ImuB form a stable complex. Gel filtration profiles of the ImuA′-ImuB complex at 14 μM (solid line) and 0.4 μM (dashed line) show identical elution volumes, indicating that the complex does not dissociate at these concentrations. For purposes of comparison, the gel filtration profiles were scaled. **B.** His-GFP-ImuB and Strep-YFP-ImuA′ were co-expressed in *E. coli* and three steps of purification were performed. The lysate was initially loaded on a His-trap column and the ImuA′-ImuB complex was eluted with an imidazole gradient (His). Next, the His-Trap elution fractions were loaded on a Strep-trap column and ImuA′-ImuB complex was eluted using a desthiobiotin gradient (Strep). Lastly, the Strep-trap's elution fraction was loaded in a size-exclusion chromatography column (SEC). Throughout all chromatography steps ImuB and ImuA′ were eluted together as a stable complex. His-GFP-ImuB and Strep-YFP-ImuA′ corresponding bands are indicated. **C.** GRS binds *M. smegmatis* DnaN. (i) Melting curve of DnaN (5 μM) in the absence (dashed line) and presence (solid line) of 15 μM GRS. Addition of GRS results in an increase of 3 °C in the protein melting temperature. (ii) Melting curve of ImuA′-ImuB complex (5 μM) in the absence (dashed line) and presence (solid line) of GRS. Addition of GRS (15 μM) does not alter the protein melting temperature (54.3°C).

**Video 1 – Time-lapse microscopy of G-ImuB and mCherry-DnaN dual reporter.** Representative time-lapse movie of the reporter strain expressing G-ImuB and mCherry-DnaN. Bacteria were imaged on fluorescence and phase channels for up to 36 hours at 10-minute intervals. Treatment with MMC (100 ng/ml) was at 0 – 4.5 hours. This experiment was repeated 6 times. Numbers indicate the hours (h) elapsed in the time-lapse experiment. 7H9, Middlebrook 7H9/OADC. MMC, Mitomycin C. Scale bar, 5 μm. G-ImuB, green; mCherry-DnaN, magenta; overlay, white.

**Video 2 – Time-lapse microscopy of V-ImuA′ and mCherry-DnaN dual reporter.** Representative time-lapse movie of the reporter strain expressing V-ImuA' and mCherry-DnaN. Bacteria were imaged on fluorescence and phase channels for up to 36 hours at 10-minute intervals. Treatment with MMC (100 ng/ml) was at 0 – 4.5 hours. This experiment was repeated 3 times. Numbers indicate the hours (h) elapsed in the time-lapse experiment. 7H9, Middlebrook 7H9/OADC. MMC, Mitomycin C. Scale bar, 5 μm. V-ImuA', green; mCherry-DnaN, magenta; overlay, white.

**Video 3 – Time-lapse microscopy of G-DnaE2 and mCherry-DnaN dual reporter.** Representative time-lapse movie of the reporter strain expressing G-DnaE2 and mCherry-DnaN. Bacteria were imaged on fluorescence and phase channels for up to 36 hours at 10-minute intervals. Treatment with MMC (100 ng/ml) was at 0 – 4.5 hours. This experiment was repeated 3 times. Numbers indicate the hours (h) elapsed in the time-lapse experiment. 7H9, Middlebrook 7H9/OADC. MMC, Mitomycin C. Scale bar, 5 μm. G-DnaE2, green; mCherry-DnaN, magenta; overlay, white.
